# The CAFA challenge reports improved protein function prediction and new functional annotations for hundreds of genes through experimental screens

**DOI:** 10.1101/653105

**Authors:** Naihui Zhou, Yuxiang Jiang, Timothy R Bergquist, Alexandra J Lee, Balint Z Kacsoh, Alex W Crocker, Kimberley A Lewis, George Georghiou, Huy N Nguyen, Md Nafiz Hamid, Larry Davis, Tunca Dogan, Volkan Atalay, Ahmet S Rifaioglu, Alperen Dalkiran, Rengul Cetin-Atalay, Chengxin Zhang, Rebecca L Hurto, Peter L Freddolino, Yang Zhang, Prajwal Bhat, Fran Supek, José M Fernández, Branislava Gemovic, Vladimir R Perovic, Radoslav S Davidović, Neven Sumonja, Nevena Veljkovic, Ehsaneddin Asgari, Mohammad RK Mofrad, Giuseppe Profiti, Castrense Savojardo, Pier Luigi Martelli, Rita Casadio, Florian Boecker, Indika Kahanda, Natalie Thurlby, Alice C McHardy, Alexandre Renaux, Rabie Saidi, Julian Gough, Alex A Freitas, Magdalena Antczak, Fabio Fabris, Mark N Wass, Jie Hou, Jianlin Cheng, Jie Hou, Zheng Wang, Alfonso E Romero, Alberto Paccanaro, Haixuan Yang, Tatyana Goldberg, Chenguang Zhao, Liisa Holm, Petri Törönen, Alan J Medlar, Elaine Zosa, Itamar Borukhov, Ilya Novikov, Angela Wilkins, Olivier Lichtarge, Po-Han Chi, Wei-Cheng Tseng, Michal Linial, Peter W Rose, Christophe Dessimoz, Vedrana Vidulin, Saso Dzeroski, Ian Sillitoe, Sayoni Das, Jonathan Gill Lees, David T Jones, Cen Wan, Domenico Cozzetto, Rui Fa, Mateo Torres, Alex Wiarwick Vesztrocy, Jose Manuel Rodriguez, Michael L Tress, Marco Frasca, Marco Notaro, Giuliano Grossi, Alessandro Petrini, Matteo Re, Giorgio Valentini, Marco Mesiti, Daniel B Roche, Jonas Reeb, David W Ritchie, Sabeur Aridhi, Seyed Ziaeddin Alborzi, Marie-Dominique Devignes, Da Chen Emily Koo, Richard Bonneau, Vladimir Gligorijević, Meet Barot, Hai Fang, Stefano Toppo, Enrico Lavezzo, Marco Falda, Michele Berselli, Silvio CE Tosatto, Marco Carraro, Damiano Piovesan, Hafeez Ur Rehman, Qizhong Mao, Shanshan Zhang, Slobodan Vucetic, Gage S Black, Dane Jo, Dallas J Larsen, Ashton R Omdahl, Luke W Sagers, Erica Suh, Jonathan B Dayton, Liam J McGuffin, Danielle A Brackenridge, Patricia C Babbitt, Jeffrey M Yunes, Paolo Fontana, Feng Zhang, Shanfeng Zhu, Ronghui You, Zihan Zhang, Suyang Dai, Shuwei Yao, Weidong Tian, Renzhi Cao, Caleb Chandler, Miguel Amezola, Devon Johnson, Jia-Ming Chang, Wen-Hung Liao, Yi-Wei Liu, Stefano Pascarelli, Yotam Frank, Robert Hoehndorf, Maxat Kulmanov, Imane Boudellioua, Gianfranco Politano, Stefano Di Carlo, Alfredo Benso, Kai Hakala, Filip Ginter, Farrokh Mehryary, Suwisa Kaewphan, Jari Björne, Hans Moen, Martti E E Tolvanen, Tapio Salakoski, Daisuke Kihara, Aashish Jain, Tomislav Šmuc, Adrian Altenhoff, Asa Ben-Hur, Burkhard Rost, Steven E Brenner, Christine A Orengo, Constance J Jeffery, Giovanni Bosco, Deborah A Hogan, Maria J Martin, Claire O’Donovan, Sean D Mooney, Casey S Greene, Predrag Radivojac, Iddo Friedberg

## Abstract

The Critical Assessment of Functional Annotation (CAFA) is an ongoing, global, community-driven effort to evaluate and improve the computational annotation of protein function. Here we report on the results of the third CAFA challenge, CAFA3, that featured an expanded analysis over the previous CAFA rounds, both in terms of volume of data analyzed and the types of analysis performed. In a novel and major new development, computational predictions and assessment goals drove some of the experimental assays, resulting in new functional annotations for more than 1000 genes. Specifically, we performed experimental whole-genome mutation screening in *Candida albicans* and *Pseudomonas aureginosa* genomes, which provided us with genome-wide experimental data for genes associated with biofilm formation and motility (*P. aureginosa* only). We further performed targeted assays on selected genes in *Drosophila melanogaster*, which we suspected of being involved in long-term memory. We conclude that, while predictions of the molecular function and biological process annotations have slightly improved over time, those of the cellular component have not. Term-centric prediction of experimental annotations remains equally challenging; although the performance of the top methods is significantly better than expectations set by baseline methods in *C. albicans* and *D. melanogaster*, it leaves considerable room and need for improvement. We finally report that the CAFA community now involves a broad range of participants with expertise in bioinformatics, biological experimentation, biocuration, and bioontologies, working together to improve functional annotation, computational function prediction, and our ability to manage big data in the era of large experimental screens.

## 1 Introduction

High-throughput nucleic acid sequencing (1) and mass-spectrometry proteomics (2) have provided us with a deluge of data for DNA, RNA, and proteins in diverse species. However, extracting detailed functional information from such data remains one of the recalcitrant challenges in the life sciences and biomedicine. Low-throughput biological experiments often provide highly informative empirical data related to various functional aspects of a gene product, but these experiments are limited by time and cost. At the same time, high-throughput experiments, while providing large amounts of data, often provide information that is not specific enough to be useful (3). For these reasons, it is important to explore computational strategies for transferring functional information from the group of functionally characterized macromolecules to others that have not been studied for particular activities (4, 5, 6, 7, 8, 9).

To address the growing gap between high-throughput data and deep biological insight, a variety of computational methods that predict protein function have been developed over the years (10, 11, 12, 13, 14, 15, 16, 17, 18, 19, 20, 21, 22, 23, 24). This explosion in the number of methods is accompanied by the need to understand how well they perform, and what improvements are needed to satisfy the needs of the life sciences community. The Critical Assessment of Functional Annotation (CAFA) is a community challenge that seeks to bridge the gap between the ever-expanding pool of molecular data and the limited resources available to understand protein function (25, 26, 27).

The first two CAFA challenges were carried out in 2010-2011 (25) and 2013-2014 (26). In CAFA1 we adopted a time-delayed evaluation method, where protein sequences that lacked experimentally verified annotations, or *targets*, were released for prediction. After the submission deadline for predictions, a subset of these targets accumulated experimental annotations over time, either as a consequence of new publications about these proteins or the biocuration work updating the annotation databases. The members of this set of proteins were used as *benchmarks* for evaluating the participating computational methods, as the function was revealed only after the prediction deadline.

CAFA2 expanded the challenge founded in CAFA1. The expansion included the number of ontologies used for predictions, the number of target and benchmark proteins, and the introduction of new assessment metrics that mitigate the problems with functional similarity calculation over concept hierarchies such as Gene Ontology (28). Importantly, we provided evidence that the top-scoring methods in CAFA2 outperformed the top scoring methods in CAFA1, highlighting that methods participating in CAFA improved over the three year period. Much of this improvement came as a consequence of novel methodologies with some effect of the expanded annotation databases (26). Both CAFA1 and CAFA2 have shown that computational methods designed to perform function prediction outperform a conventional function transfer through sequence similarity (25, 26).

In CAFA3 (2016-2017) we continued with all types of evaluations from the first two challenges and additionally performed experimental screens to identify genes associated with specific functions. This allowed us to provide unbiased evaluation of the term-centric performance based on a unique set of benchmarks obtained by assaying *Candida albicans, Pseudomonas aeruginosa* and *Drosophila melanogaster*. We also held a challenge following CAFA3, dubbed CAFA-*π*, to provide the participating teams another opportunity to develop or modify prediction models. The genome-wide screens on *C. albicans* identified 240 genes previously not known to be involved in biofilm formation, whereas the screens on *P. aeruginosa* identified 532 new genes involved in biofilm formation and 403 genes involved in motility. Finally, we used CAFA predictions to select genes from *D. melanogaster* and assay them for long-term memory involvement. This experiment allowed us to both evaluate prediction methods and identify eleven new fly genes involved in this biological process (29). Here we present the outcomes of the CAFA3 challenge, as well as the accompanying challenge CAFA-*π*, and discusses further directions for the community interested in the function of biological macromolecules.

## 2 Results

### 2.1 Top methods have slightly improved since CAFA2

One of CAFA’s major goals is to quantify the progress in function prediction over time. We therefore conducted comparative evaluation of top CAFA1, CAFA2, and CAFA3 methods according to their ability to predict Gene Ontology (28) terms on a set of common benchmark proteins. This benchmark set was created as an intersection of CAFA3 benchmarks (proteins that gained experimental annotation after the CAFA3 prediction submission deadline), and CAFA1 and CAFA2 target proteins. Overall, this set contained 377 protein sequences with annotations in the Molecular Function Ontology (MFO), 717 sequences in the Biological Process Ontology (BPO) and 548 sequences in the Cellular Component Ontology (CCO), which allowed for a direct comparison of all methods that have participated in the challenges so far. The head-to-head comparisons in MFO, BPO, and CCO between top five CAFA3 and CAFA2 methods are shown in Figure 1. CAFA3 and CAFA1 comparisons are shown in Figure S1 in the Supplemental Materials.

**Figure 1:**
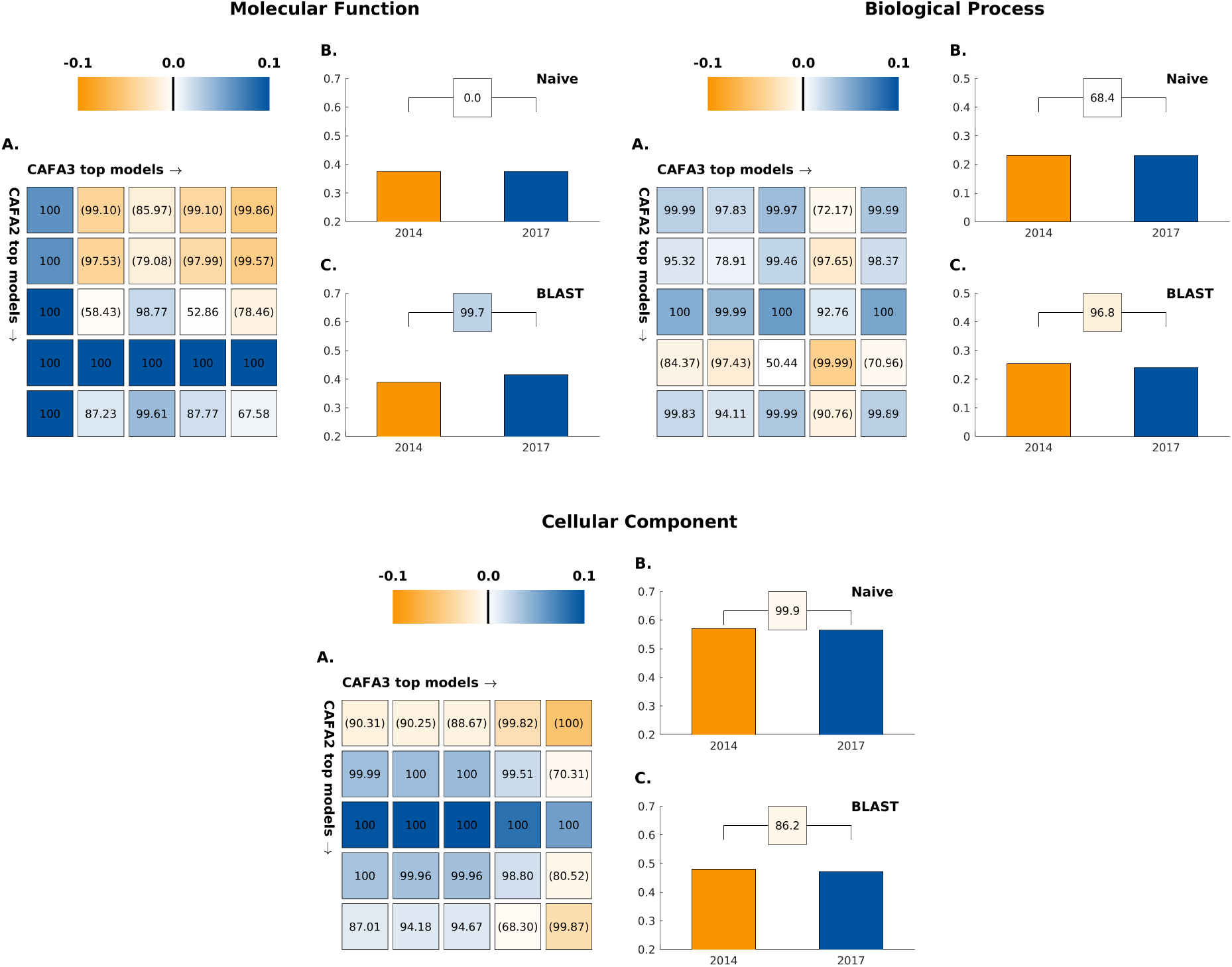
A comparison in *F*_max_ between the top-five CAFA2 models against the top-five CAFA3 models. Colored boxes encode the results such that (1) the colors indicate margins of a CAFA3 method over a CAFA2 method in *F*_max_ and (2) the numbers in the box indicate the percentage of wins. A: CAFA2 top-five models (rows, from top to bottom) against CAFA3 top-five models (columns, from left to right). B: Comparison of performance (*F*_max_) of Naïve baselines trained respectively on SwissProt2014 and SwissProt2017. C: Comparison of performance (*F*_max_) of BLAST baselines trained on SwissProt2014 and SwissProt2017. Statistical significance was assessed using 10,000 bootstrap samples of benchmark proteins.

We first observe that, in effect, the performance of baseline methods (25, 26) has not improved since CAFA2. The Naïve method, which uses the term frequency in the existing annotation database as prediction score for every input protein, has the same *F*_max_ performance using both annotation database in 2014 (when CAFA2 was held) and in 2017 (when CAFA3 was held), which suggests little change in term frequencies in the annotation database since 2014. On the other hand, BLAST-based annotation transfer, tells a contrasting tale between ontologies. In MFO, the BLAST method based on the existing annotations in 2017 is slightly but significantly better than the BLAST method based on 2014 training data. In BPO and CCO, however, the BLAST based on the later database has not outperformed its earlier counterpart, although the changes in effect size (absolute change in *F*_max_) in both ontologies are small.

When surveying all three CAFA challenges, the performance of both baseline methods has been relatively stable, with some fluctuations of BLAST. Such performance of direct sequence-based function transfer is surprising, given the steady growth of annotations in UniProt-GOA (30); i.e., there were 259,785 experimental annotations in 2011, 341,938 in 2014 and 434,973 in 2017, but there does not seem to be a definitive trend with the BLAST method, as they go up and down in *F*_max_ across ontologies. We conclude from these observations on the baseline methods that first, the ontologies are in different annotation states and should not be treated as a whole. Second, methods that perform direct function transfer based on sequence similarity do not necessarily benefit from a larger training dataset. Although the performance observed in our work is also dependent on the benchmark set, it appears that the annotation databases remain sparsely populated to effectively exploit function transfer by sequence similarity, thus justifying the need for advanced methodology development for this problem.

Head-to-head comparisons of the top five CAFA3 methods against top five CAFA2 methods show mixed results. In MFO, the top CAFA3 method, GOLabeler (23) outperformed all CAFA2 methods by a considerable margin, as shown in Figure 2. The rest of the four CAFA3 top methods did not perform as well as the top two methods of CAFA2, although only to a limited extent, with little change in *F*_max_. Of the top 12 methods ranked in MFO, seven are from CAFA3, five are from CAFA2 and none are from CAFA1. Despite the increase in database size, the majority of function prediction methods do not seem to have improved in predicting protein function in MFO since 2014, except for one method that stood out. In BPO, the top three methods in CAFA3 outperformed their CAFA2 counterparts, but with very small margins. Out of the top 12 methods in BPO, eight are from CAFA3, four are from CAFA2 and none are from CAFA1. Finally, in CCO, although 8 out of top 12 methods over all CAFA challenges come from CAFA3, the top method is from CAFA2. The differences between the top performing methods are small, as in the case of BPO.

**Figure 2:**
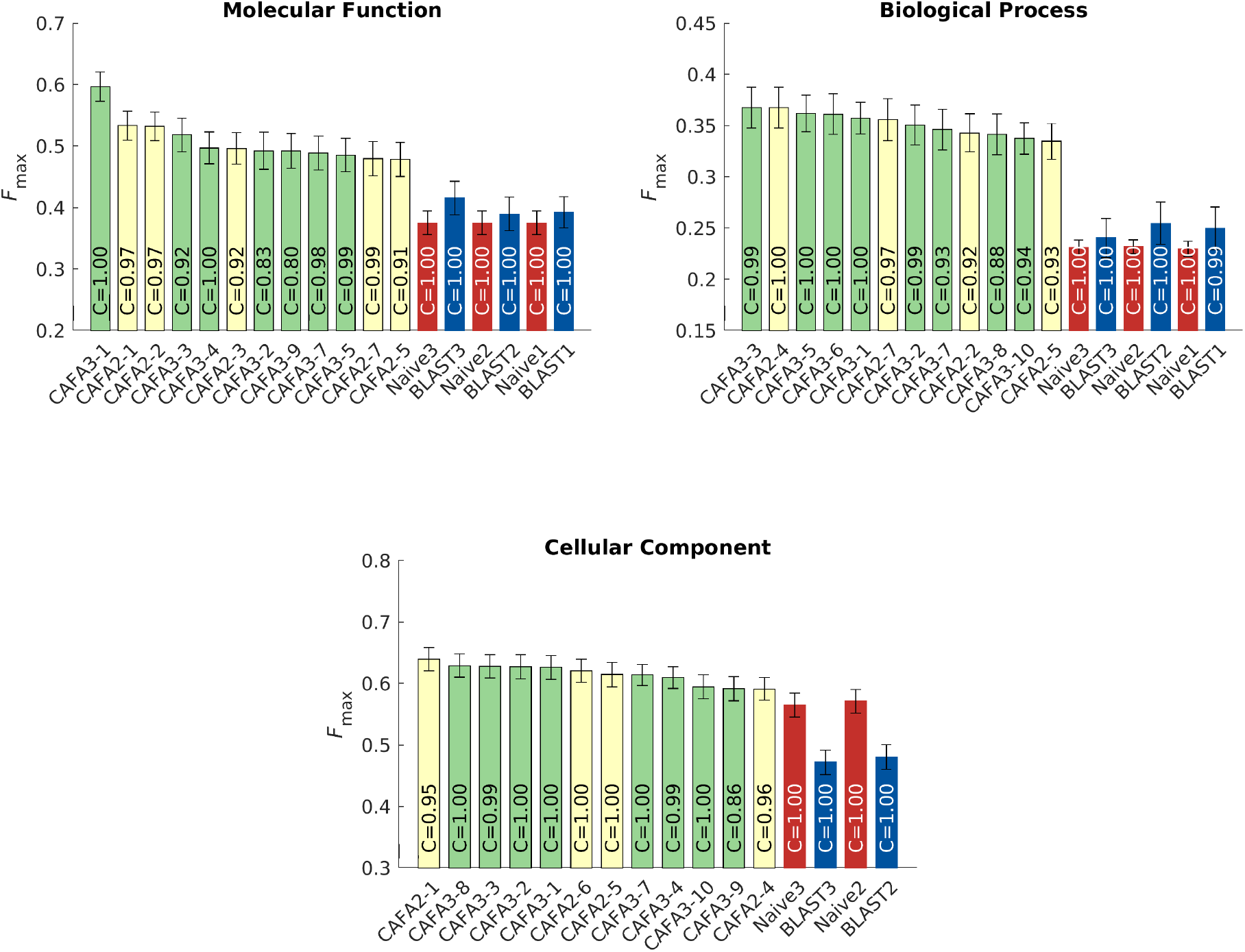
Performance evaluation based on *F*_max_ for top CAFA1, CAFA2 and CAFA3 methods. The top 12 methods are shown in this barplot ranked in descending order fro left to right. The baseline methods are appended to the right, they were trained on training data from 2017, 2014 and 2011 respectively. Coverage of the methods were shown as text inside the bars. Coverage is defined as percentage of proteins in the benchmark that are predicted by the methods. Color scheme: CAFA2: ivory; CAFA3: green; Naïve: red; BLAST: blue. Note that in MFO and BPO, CAFA1 methods were ranked but not displayed. CAFA1 challenge did not collect predictions for CCO.

The performance of top methods in CAFA2 was significantly better than of those in CAFA1, and it is interesting to note that this trend has not continued in CAFA3. This could be due to many reasons, such as the quality of the benchmark sets, the overall quality of the annotation database, the quality of ontologies or a relatively short period of time between challenges.

### 2.2 Protein-centric evaluation

The *protein-centric* evaluation measures the accuracy of assigning GO terms to a protein. This performance is shown in Figures 3, 4 and 5.

**Figure 3:**
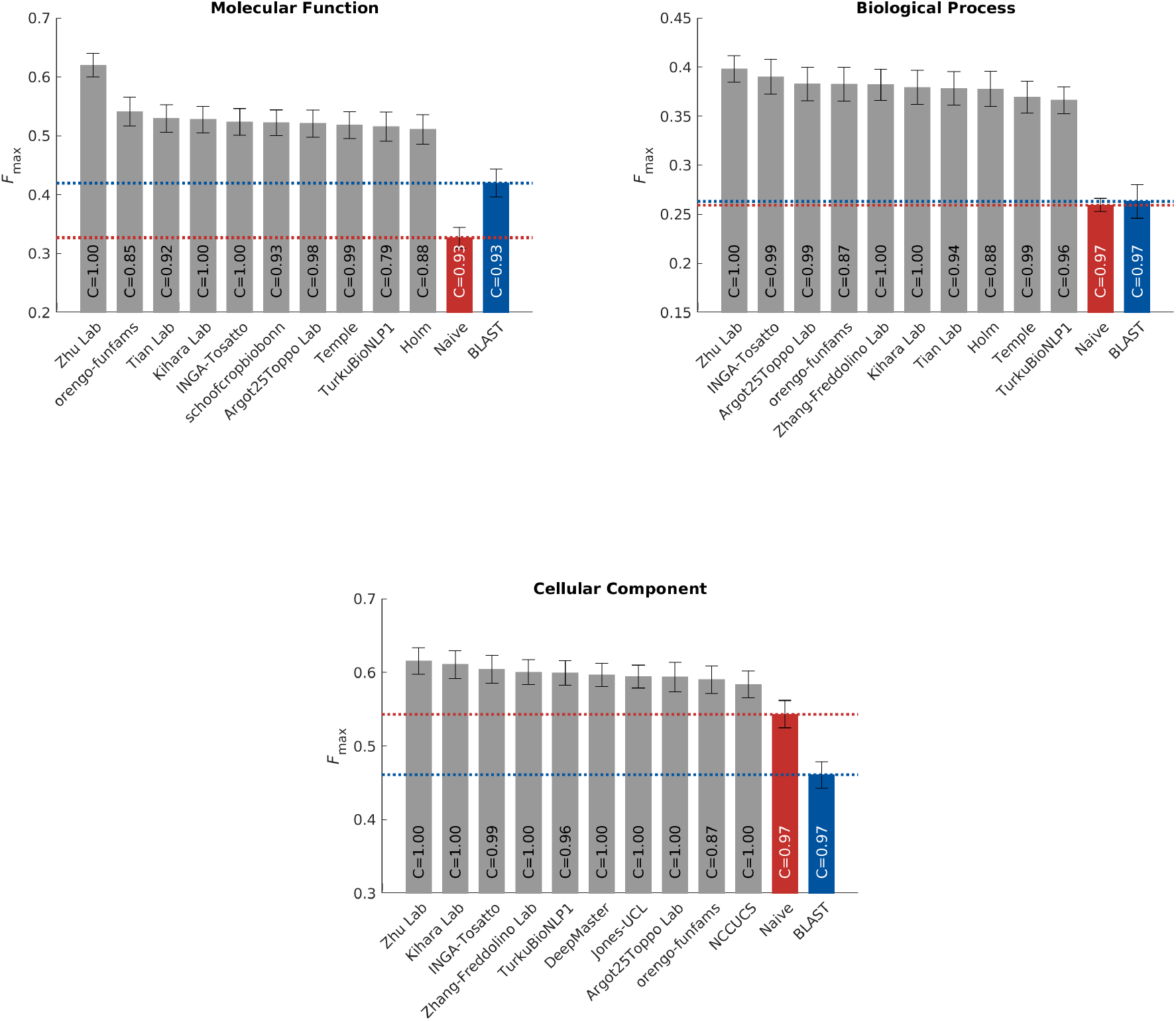
Performance evaluation based on *F*_max_ for the top-performing methods in three ontologies. The 95% confidence interval was estimated using 10,000× bootstrap on the benchmark set. Coverage of the methods were shown as text inside the bars. Coverage is defined as percentage of proteins in the benchmark that are predicted by the methods.

**Figure 4:**
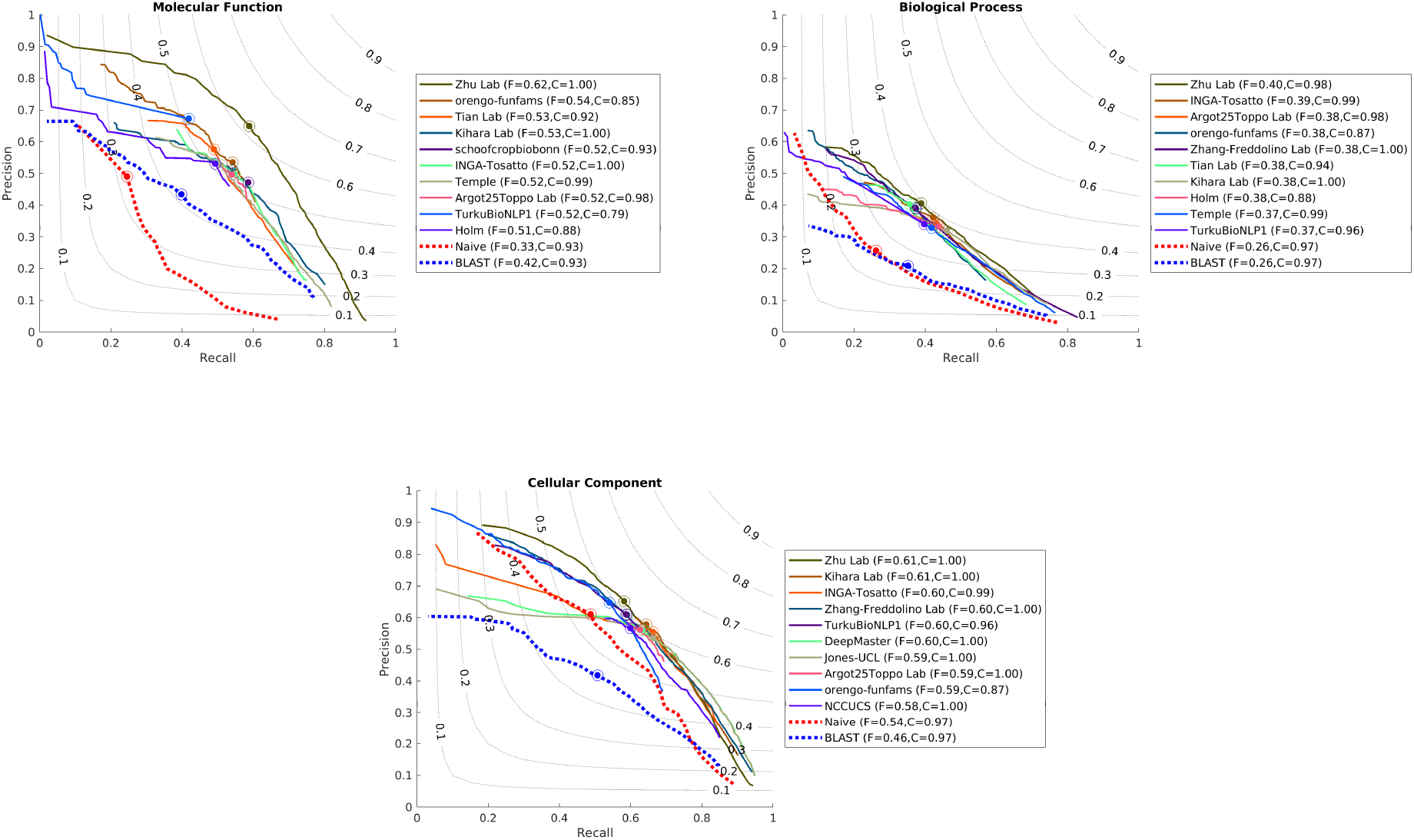
Precision Recall curves for the top-performing methods

**Figure 5:**
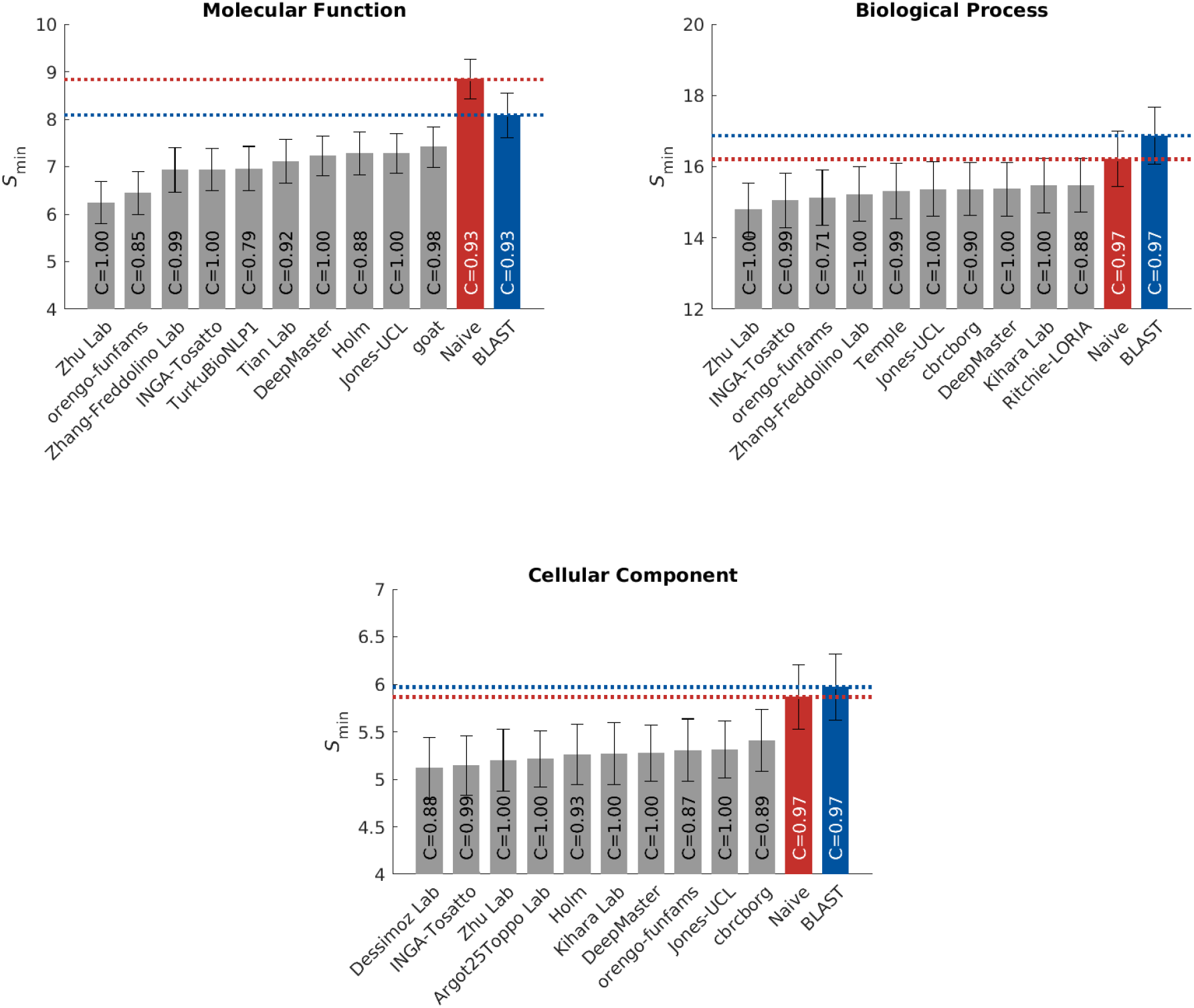
Evaluation based on the *S*_min_ for the top-performing methods

We observe that all top methods outperform the baselines with the patterns of performance consistent with CAFA1 and CAFA2 findings. Predictions of MFO terms achieved the highest *F*_max_ compared with predictions in the other two ontologies. BLAST outperforms Naïve in predictions in MFO, but not in BPO or CCO. This is because sequence similarity based methods such as BLAST tend to perform best when transferring basic biochemical annotations such as enzymatic activity. Functions in biological process, such as pathways, may not be as preserved by sequence similarity, hence the poor BLAST performance in BPO. The reasons behind the difference among the three ontologies include the structure and complexity of the ontology as well as the state of the annotation database, as discussed previously (26, 31). It is less clear why the performance in CCO is weak, although it might be hypothesized that such performance is related to the structure of the ontology itself (31).

The top performing method in MFO did not have as high an advantage over others when evaluated using the *S*_min_ metric. The *S*_min_ metric weights GO terms by conditional information content, since the prediction of more informative terms are more desirable than less informative, more general, terms. This could potentially explain the smaller gap between the top predictor and the rest of the pack in *S*_min_. The weighted *F*_max_ and normalized *S*_min_ evaluations can be found in Figures S4 and S5.

### 2.3 Species-specific categories

The benchmarks in each species were evaluated individually as long as there were at least 15 proteins per species. Here we present results on both eukaryotic and prokaryotic species (Figure 6). We observed that different methods could perform differently on different species. As shown in Figure 14, bacterial proteins make up a small portion of all benchmark sequences, so their effects on the performances of the methods are often masked. Species-specific analyses are thus meaningful to researchers studying certain organisms. Evaluation results on individual species including human (Figure S6), *Arabidopsis thaliana* (Figure S7) and *Escherichia coli* (Figure S10) can be found in Supplemental Materials (Figures S6-S14).

**Figure 6:**
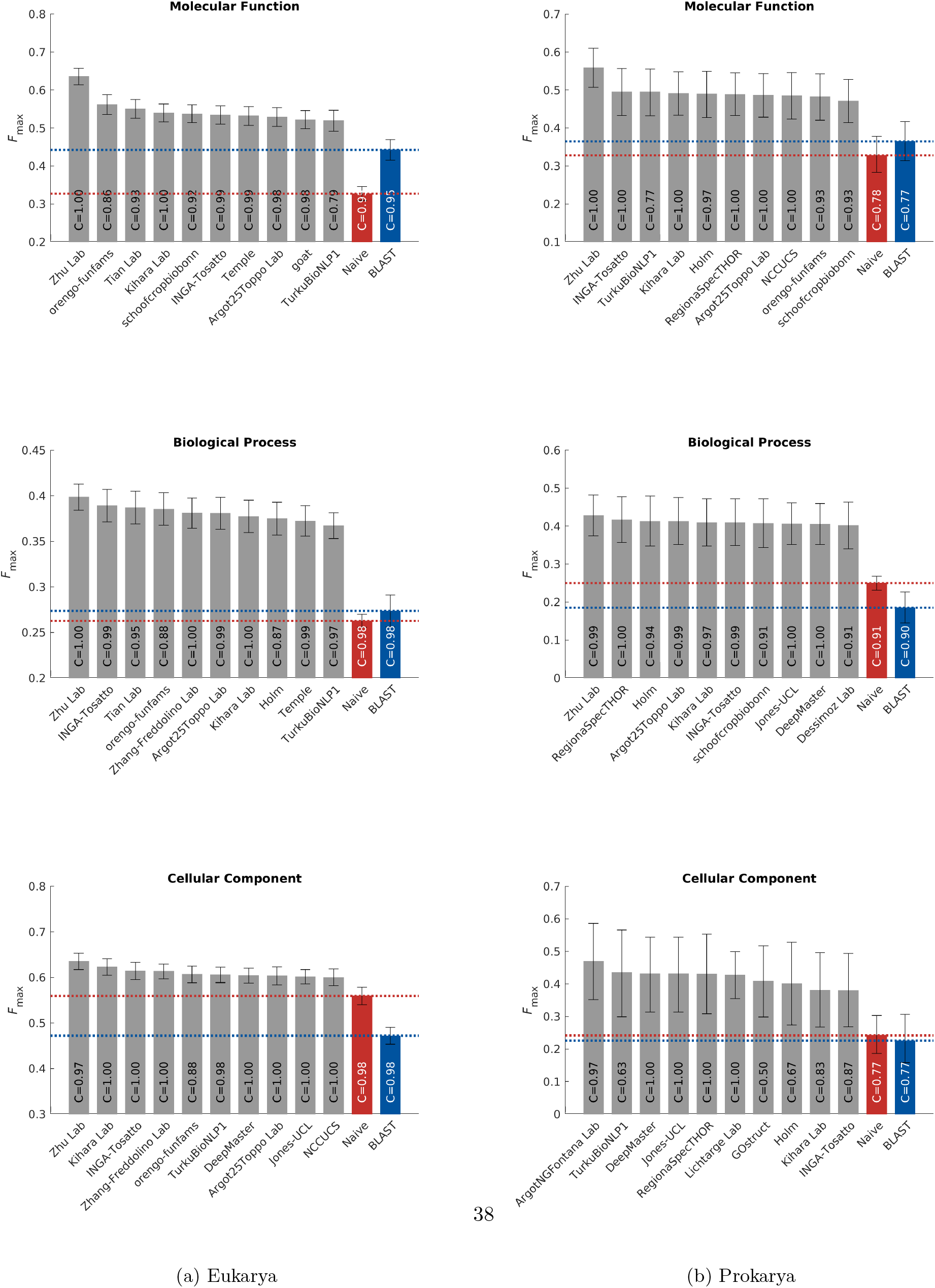
Evaluation based on the *F*_max_ for the top-performing methods in eukaryotic and prokaryotic species

### 2.4 Diversity of methods

It was suggested in the analysis of CAFA2 that ensemble methods that integrate data from different sources have the potential of improving prediction accuracy (32). Multiple data sources, including sequence, structure, expression profile and so on are all potentially predictive of the function of the protein. Therefore, methods that take advantage of these rich sources as well as existing techniques from other research groups might see improved performance. Indeed, the one method that stood out from the rest in CAFA3 and performed significantly better than all methods across three challenges, is a machine learning based ensemble method (23). Therefore, it is important to analyze what information sources and prediction algorithms are better at predicting function. Moreover, the similarity of the methods might explain the limited improvement in the rest of the methods in CAFA3.

The top CAFA2 and CAFA3 methods are very similar in performance, but that could be a result of aggregating predictions of different proteins to one metric. When computing the similarity of each pair of methods as the reciprocal of the Euclidean distance of prediction scores (Figure 7), we are not interested whether these predictions are correct according to the benchmarks, but simply whether they are similar to one another. Top CAFA2 and CAFA3 methods are more similar than with CAFA1 models. It is clear that some top methods are heavily based on the Naïve and BLAST baseline methods. It is interesting to note that the top two best methods in BPO are not similar to any other top methods. The same pattern was observed for CAFA2 methods.

**Figure 7:**
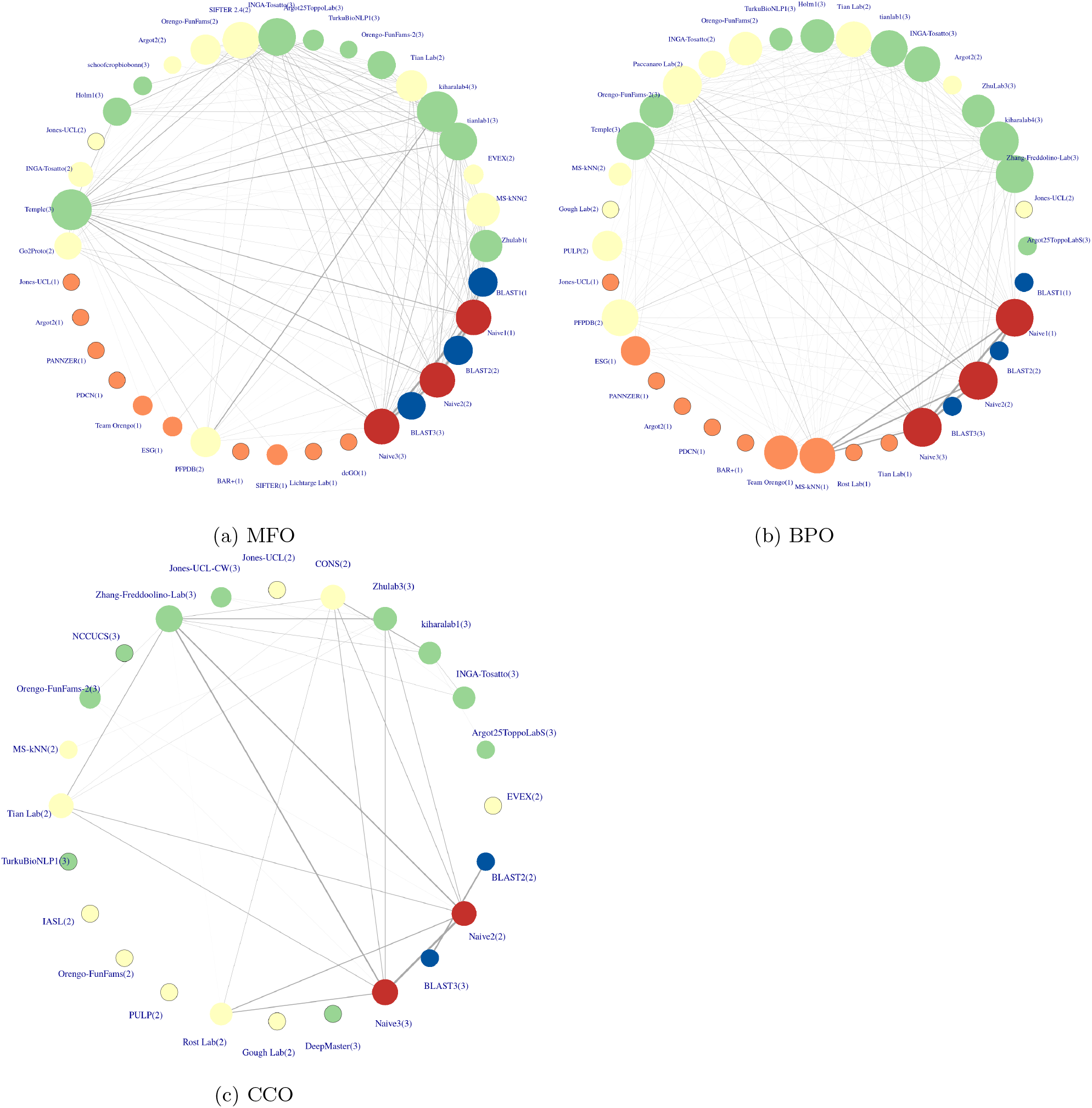
Similarity networks of top 10 methods from CAFA1, CAFA2 and CAFA3. The team names are displayed together with which CAFA challenge they come from in parenthesis. Similarity is calculated as the reciprocal of the Euclidean distance of the prediction scores from each pair of methods. A 0.07 cutoff was applied to the Euclidean distances, i.e. an edge exists if the Euclidean distance is lower than the cutoff. Edge width is directly proportional to similarity, except at the three edges between the three Naïve methods, where the similarity is much larger than the rest. Vertex size is directly proportional to number of edges, or degree of a vertex. Singletons, or vertices without any edges are framed with black circles. The nodes are ranked counter-clockwise, starting after ‘BLAST1’, by *F*_max_ performance in the intersection set of benchmarks in Section 2.1. Color scheme: CAFA1: orange; CAFA2: ivory; CAFA3: green; Naïve: red; BLAST: blue.

Participating teams also provided keywords that describe their approach to function prediction with their submissions. A list of keywords was given to the participants, listed in Page 24 of Supplementary Materials. Figure 8 shows the frequency of each keyword. In addition, we have weighted the frequency of the keywords with the prediction accuracy of the specific method. Machine learning and sequence alignment remain the most-used approach by scientists predicting in all three ontologies. By raw count, machine learning is more popular than sequence alignment, but once adjusted by performance, they are almost identical. This indicates that methods that use sequence alignments are more helpful in predicting the correct function than the popularity of their use suggests.

**Figure 8:**
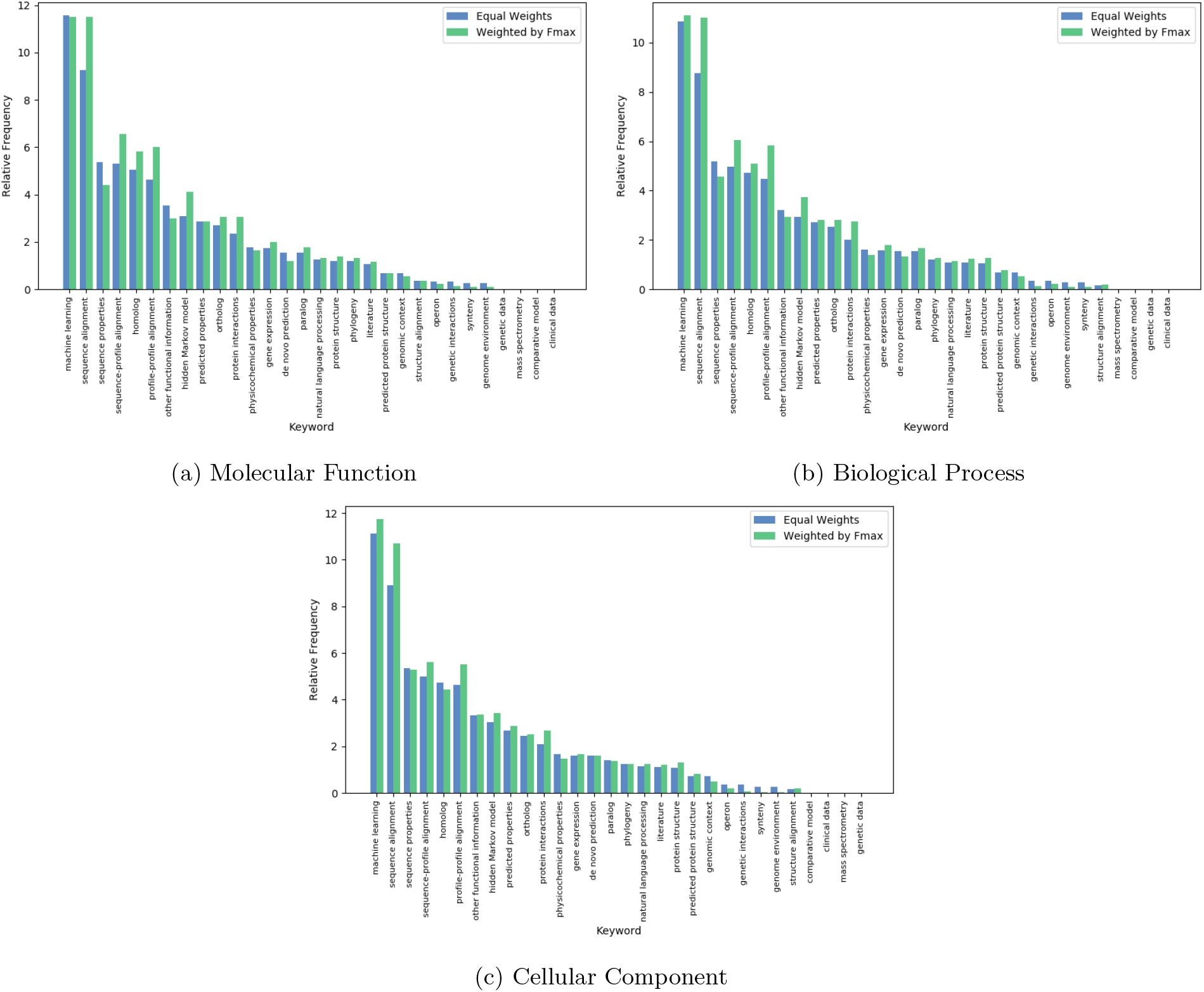
Keyword analysis of all CAFA3 participating methods. Both relative frequency of the keywords and weighted frequency are provided. The weighted frequencies accounts for the performance of the the particular model using the given keyword. If that model performs well (with high *F*_max_) then it gives more weight to the calculation of the total weighted average of that keyword.

### 2.5 Evaluation via molecular screening

Databases with proteins annotated by biocuration, such as UniProt knowledge base, have been the primary source of benchmarks in the CAFA challenges. New to CAFA3, we also evaluated the extent to which methods participating in CAFA could predict the results of genetic screens in model organisms done specifically for this project. Predicting GO terms for a protein (protein-centric) and predicting which proteins are associated with a given function (term-centric) are related but different computational problems: the former is a multi-label classification problem with a structured output, while the latter is a binary classification task. Predicting the results of a genome-wide screen for a single or a small number of functions fits the term-centric formulation. To see how well all participating CAFA methods perform term-centric predictions, we mapped results from the protein-centric CAFA3 methods onto these terms. In addition we held a separate CAFA challenge, CAFA-*π* whose purpose was to attract additional submissions from algorithms that specialize in term-centric tasks.

We performed screens for three functions in three species, which we then used to assess protein function prediction. In the bacterium *Pseudomonas aeruginosa* and the fungus *Candida albicans* we performed genome-wide screens capable of uncovering genes with two functions, biofilm formation (GO:0042710) and motility (for *P. aeruginosa* only) (GO:0001539), as described in Methods. In *Drosophila melanogaster* we performed targeted assays, guided by previous CAFA submissions, of a selected set of genes and assessed whether or not they affected long-term memory (GO:0007616).

We discuss the prediction results for each function below in detail. The performance, as assessed by the genome-wide screens, was generally lower than in the protein-centric evaluations that were curation driven. We hypothesize that it may simply be more difficult to perform term-centric prediction for broad activities such as biofilm formation and motility. For *P. aeruginosa*, an existing compendium of gene expression data was already available (33). We used the Pearson correlation over this collection of data to provide a complementary baseline to the standard BLAST approach used throughout CAFA. We found that an expression-based method outperformed the CAFA participants, suggesting that success on certain term-centric challenges will require the use of different types of data. On the other hand, the performance of the methods in predicting long-term memory in the Drosophila genome was relatively accurate.

#### 2.5.1 Biofilm formation

In March 2018, there were 3019 annotations to biofilm formation (GO:0042710) and its descendent terms across all species, of which 325 used experimental evidence codes. These experimentally annotated proteins included 131 from the Candida Genome Database (34) for *C. albicans* and 29 for *P. aeruginosa*, the two organisms that we screened.

Of the 2746 genes we screened in the *Candida albicans* colony biofilm assay, 245 were required for the formation of wrinkled colony biofilm formation (Table 1). Of these, only five were already annotated in UniProt: *MOB, EED1* (*DEF1*), and *YAK1*, which encode proteins involved in hyphal growth, an important trait for biofilm formation (35, 36, 37, 38). Also, *NUP85*, a nuclear pore protein involved in early phase arrest of biofilm formation (39) and *VPS1*, which contributes to protease secretion, filamentation, and biofilm formation (40). Of the 2063 proteins that we did not find to be associated with biofilm formation, 29 were annotated to the term in the GOA database. Some of the proteins in this category highlight the need for additional information to GO term annotation. For example, Wor1 and the pheromone receptor are key for biofilm formation in strains under conditions in which the mating pheromone is produced (41), but not required in the monocultures of the commonly studied a/*α* mating type strain used here.

**Table 1:**
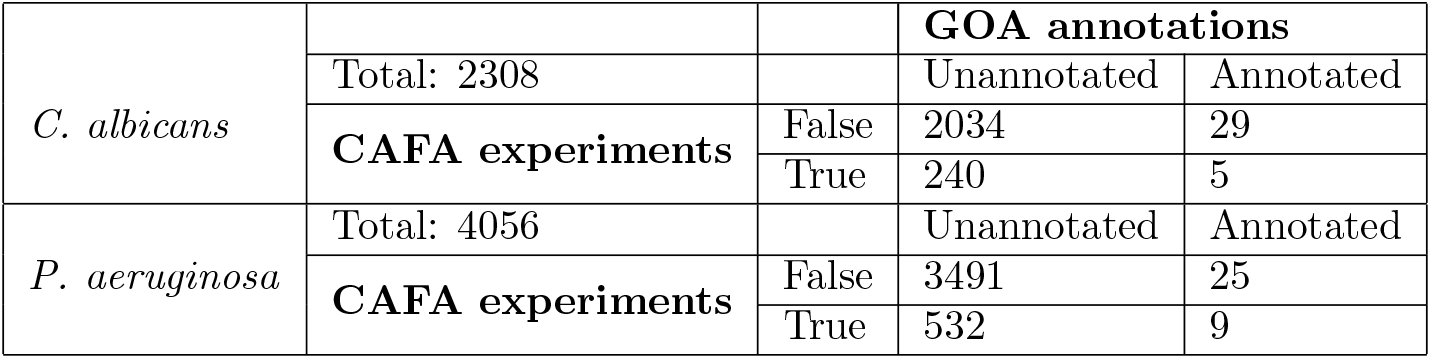
Number of proteins in Candida albicans and *Pseudomonas aeruginosa* associated with function Biofilm formation (GO:0042710) in the GOA databases versus experimental results.

|  |  |  | GOA annotations |  |
| --- | --- | --- | --- | --- |
|  | Total: 2308 |  | Unannotated | Annotated |
| *C. albicans* | CAFA experiments | False | 2034 | 29 |
|  |  | True | 240 | 5 |
| *P. aeruginosa* | Total: 4056 |  | Unannotated | Annotated |
|  | CAFA experiments | False | 3491 | 25 |
|  |  | True | 532 | 9 |

No method in CAFA-*π* or CAFA3 (not shown) exceeded an AUC of 0.60 on this term-centric challenge (Figure 9) for either species. Performance for the best methods slightly exceeded a BLAST-based baselines. In the past, we have found that predicting BPO terms, such as biofilm formation, resulted in poorer method performance than predicting MFO terms. Many CAFA methods use sequence alignment as their primary source of information (Section 2.4). For *Pseudomonas aeruginosa* a pre-built expression compendium was available from prior work (33). Where the compendium was available, simple gene-expression based baselines were the best performing approaches. This suggests that successful term-centric prediction of biological processes may need to rely more heavily on information that is not sequence-based, and, as previously reported, may require methods that use broad collections of gene expression data (42, 43).

**Figure 9:**
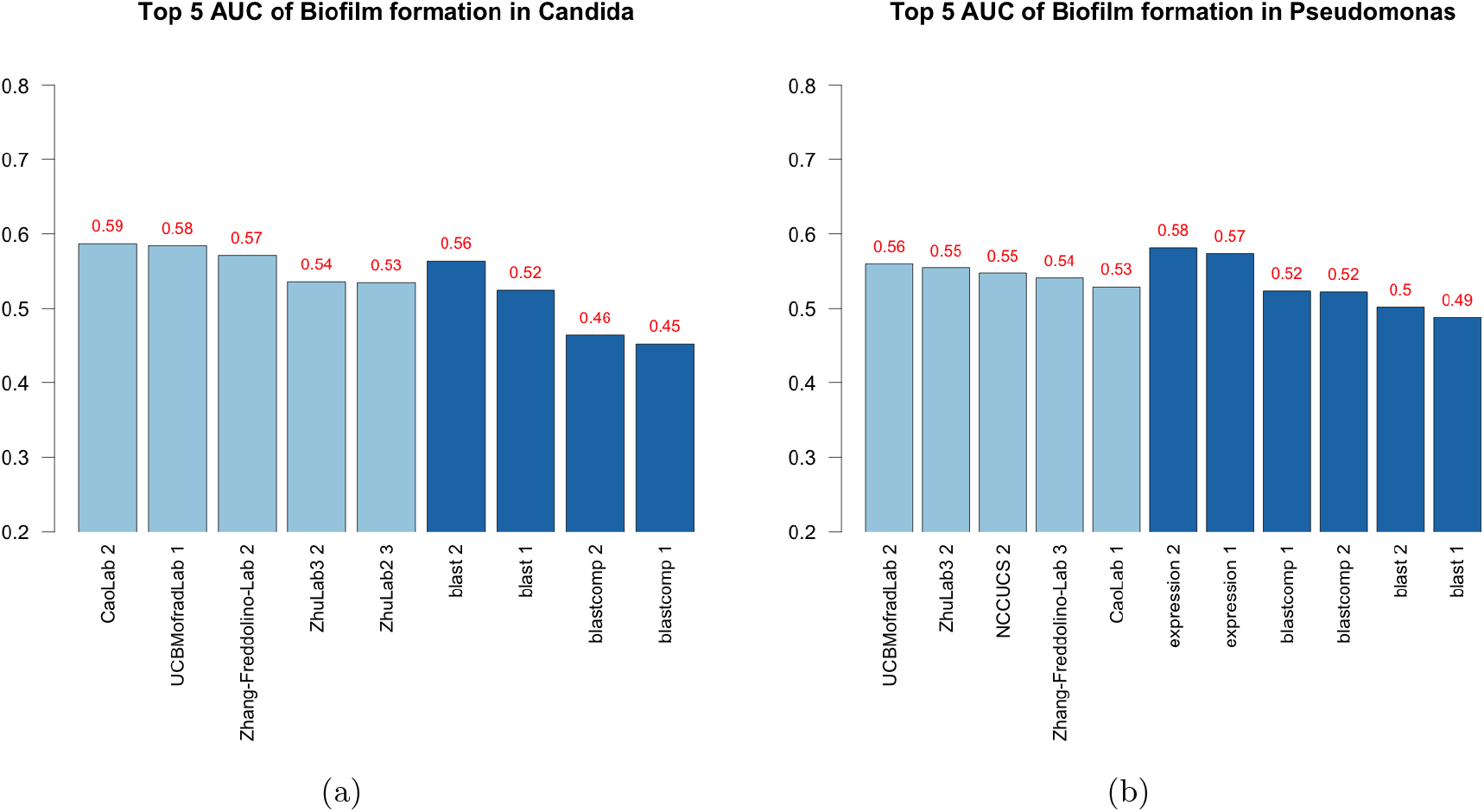
AUROC of top 5 teams in CAFA-*π*. The best performing model from each team is picked for the top five teams, regardless of whether that model is submitted as model 1. Four baseline models all based on BLAST were computed for *Candida*, while six baseline models were computed for *Pseudomonas*, including two based on Expression profiles.

**Figure 10:**
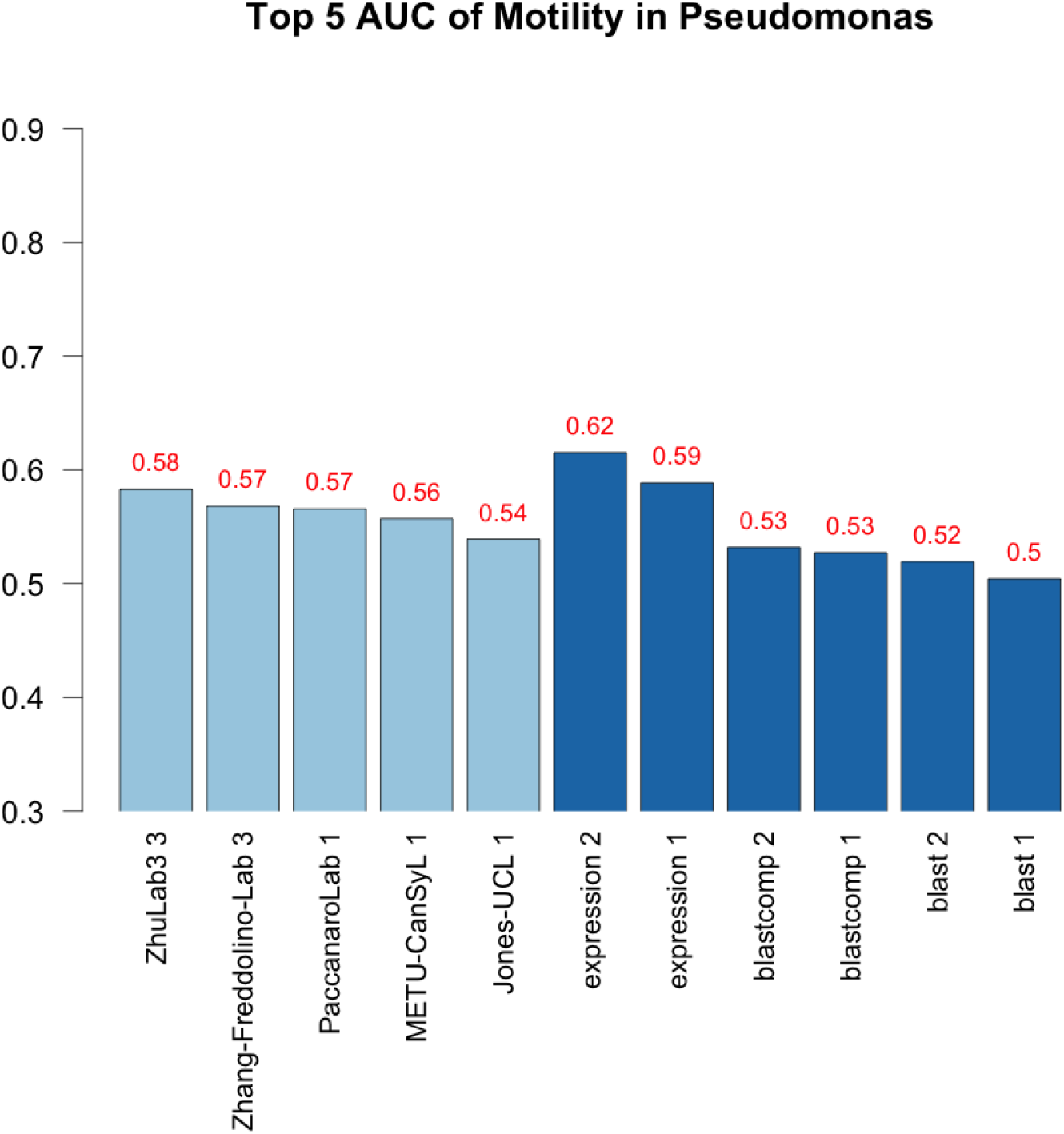
AUROC of top 5 teams in CAFA-*π*. The best performing model from each team is picked for the top five teams, regardless of whether that model is submitted as model 1.

#### 2.5.2 Motility

In March 2018 there were 302,121 annotations for proteins with the GO term: cilium or flagellum-dependent cell motility (GO:0001539) and its descendent terms, which included cell motility in all eukaryotic (GO:0060285), bacterial (GO:0071973) and archael (GO:0097590) organisms. Of these, 187 had experimental evidence codes and the most common organism to have annotations was *P. aeruginosa*, on which our screen was performed (Table S2).

As expected, mutants defective in the flagellum or its motor were defective in motility (*fliC* and other *fli* and *flg* genes). For some of the genes that were expected, but not detected, the annotation was based on experiments performed in a medium different from what was used in these assays. For example, PhoB regulates motility but only when phosphate concentration is low (44). Among the genes that were scored as defective in motility, some are known to have decreased motility due to over production of carbohydrate matrix material (*bifA*) (45), or the absence of directional swimming due to absence of chemotaxis functions (e.g., *cheW, cheA*) and others likely showed this phenotype because of a medium specific requirement such as biotin (*bioA, bioC*, and *bioD*) (46). Table 2 shows the contingency table for number of proteins that are detected by our experiment versus GOA annotations.

**Table 2:** Number of proteins in *Pseudomonas aeruginosa* associated with function Motility (GO:0001539) in the GOA databases versus experimental results.

| Total: 3630 |  | GOA annotations |  |
| --- | --- | --- | --- |
|  |  | Unannotated | Annotated |
| CAFA experiments | False | 3195 | 12 |
|  | True | 403 | 21 |

The results from this evaluation were consistent with what we observed for biofilm formation. Many of the genes annotated as being involved in biofilm formation were identified in the screen. Others that were annotated as being involved in biofilm formation did not show up in the screen because the strain background used here, strain PA14, uses the exoploysaccharide matrix carbohydrate Pel (47) in contrast to the Psl carbohydrate used by another well characterized strain, strain PAO1 (48, 49). The *psl* genes were known to be dispensable for biofilm formation in the strain PA14 background and this nuance highlights the need for more information to be taken into account when making predictions.

The CAFA-*π* methods outperformed our BLAST-based baselines but failed to outperform expression-based baselines. Transferred methods from CAFA3 also did not outperform these baselines. It is important to note this consistency across terms, reinforcing the finding that term-centric prediction of biological processes is likely to require non-sequence information to be included.

#### 2.5.3 Long-term memory in *D. melanogaster*

Prior to our experiments, there were 1901 annotations made in long-term memory, including 283 experimental annotations. *Drosophila melanogaster* had the most annotated proteins of long-term memory with 217, while human has 7, as shown in Table S3.

We performed RNAi experiments in *Drosophila melanogaster* to assess whether 29 target genes were associated with long-term memory (GO:0007616); for details on target selection, see (29). None of the 29 genes had an existing annotation in the GOA database. Because no genome-wide screen results were available, we did not release this as part of CAFA-*π* and instead relied only on the transfer of methods that predicted “long-term memory” at least once in *D. melanogaster* from CAFA3. Results from this assessment were more promising than our findings from the genome-wide screens in microbes (Figure 11). Certain methods performed well, substantially exceeding the baselines.

**Figure 11:**
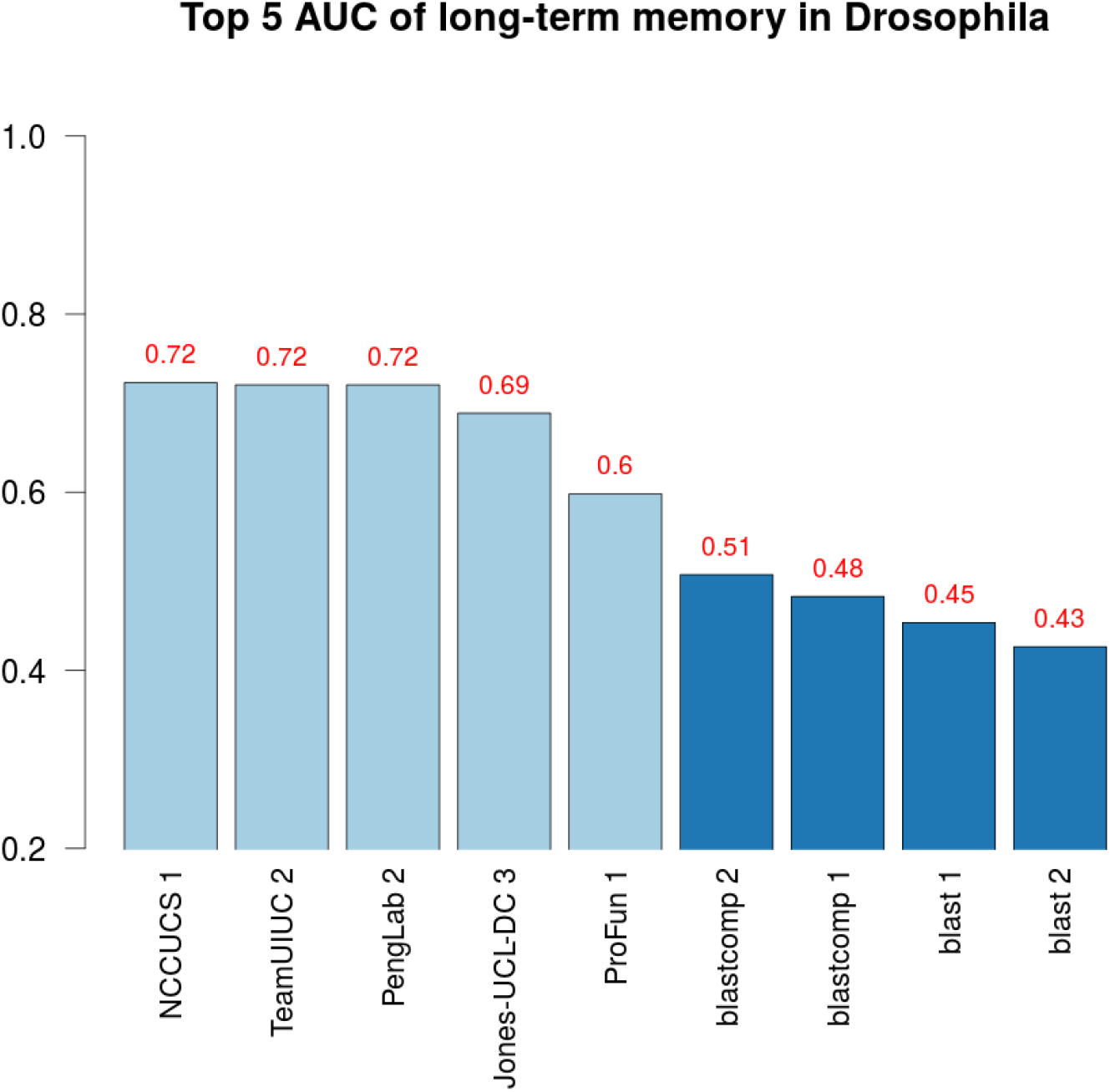
AUROC of top five teams in CAFA3. The best performing model from each team is picked for the top five teams, regardless of whether that model is submitted as model 1.

### 2.6 Participation Growth

The CAFA challenge has seen growth in participation, as shown in Figure 12. To cope with the increasingly large data size, CAFA3 utilized the Synapse (50) online platform for submission. Synapse allowed for easier access for participants, as well as easier data collection for the organizers. The results were also released to the individual teams via this online platform. During the submission process, the online platform also allows for customized format checkers to ensure the quality of the submission.

**Figure 12:**
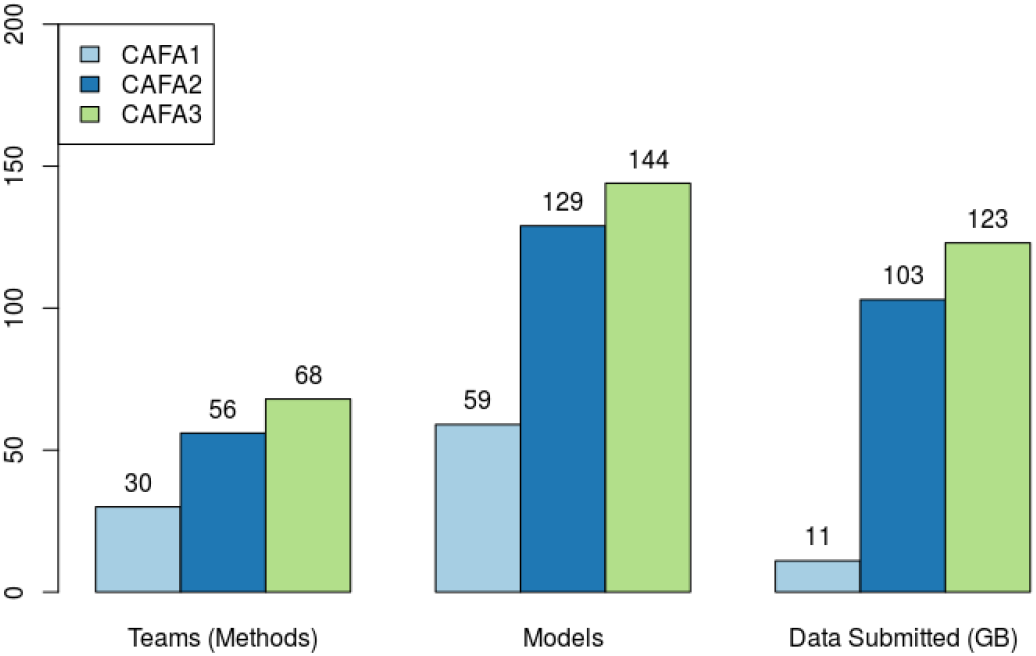
CAFA participation has been growing. Each Principle Investigator is allowed to head multiple teams, but each member can only belong to one team. Each team can submit up to three models.

## 3 Methods

### 3.1 Benchmark collection

In CAFA3, we adopted the same benchmark generation methods as CAFA1 and CAFA2, with a similar time-line (Figure 13). The crux of a time-delayed challenge is the annotation growth period between time *t*_0_ and *t*_1_. All target proteins that have gained experimental annotation during this period are taken as benchmarks in all three ontologies. “No-knowledge” (NK, no prior experimental annotations) and “Limited-knowledge” (LK, partial prior experimental annotations) benchmarks were also distinguished based on whether the newly-gained experimental annotation is in an ontology that already have experimental annotations or not. Evaluation results in Figures 3, 4, and 5 are made using the No-knowledge benchmarks. Evaluation results on the Limited-knowledge benchmarks are shown in Figure S3 in the Supplemental Materials. For more information regarding NK and LK designations, please refer to the Supplemental Materials and the CAFA2 paper (26).

**Figure 13:**
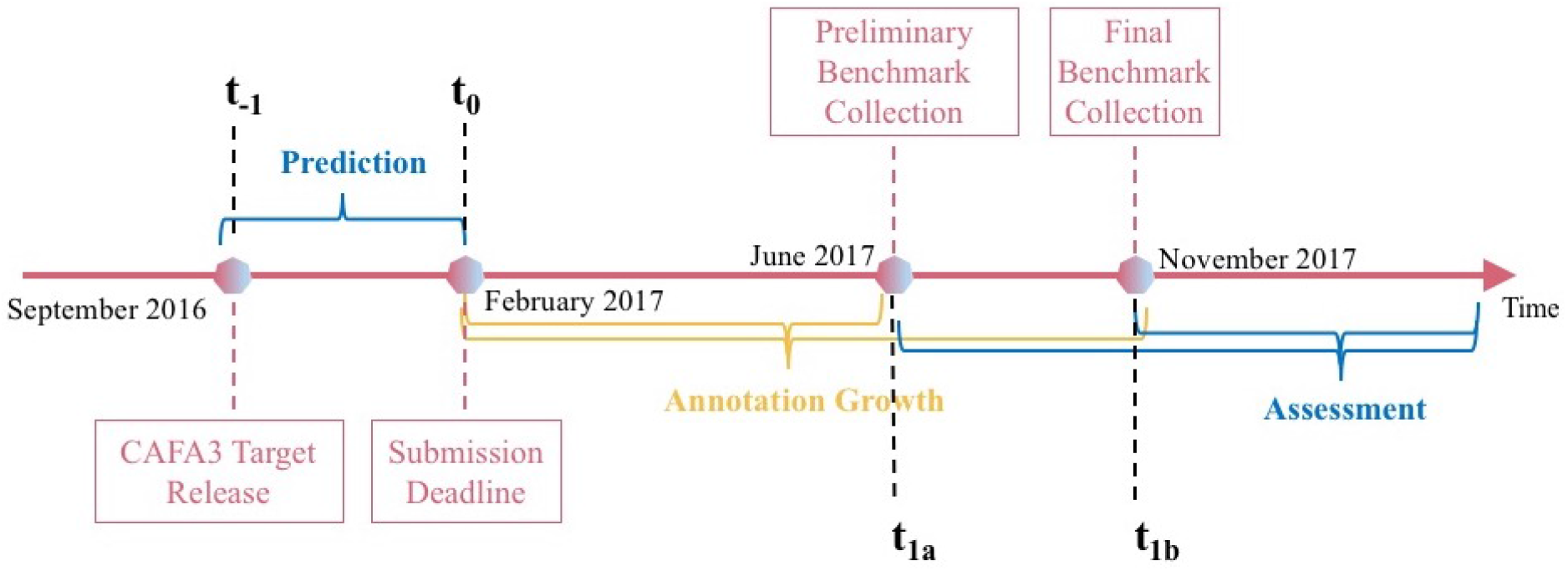
CAFA3 timeline

After collecting these benchmarks, we performed two major deletions from the benchmark data. Upon inspecting the taxonomic distribution of the benchmarks, we noticed a large number of new experimental annotations from *Candida albicans*. After consulting with UniProt-GOA, we determined these annotations have already existed in the Candida Genome Database long before 2018, but were only recently migrated to GOA. Since these annotations were already in the public domain before the CAFA3 submission deadline, we have deleted any annotation from *Candida albicans* with an assigned date prior to our CAFA3 submission deadline. Another major change is the deletion of any proteins with only a protein-binding (GO:0005515) annotation. Protein-binding is a highly generalized function description, does not provide more specific information about the actual function of a protein, and in many cases may indicate a non-functional, non-specific binding. If it is the only annotation that a protein has gained, then it is hardly an advance in our understanding of that protein, therefore we deleted these annotations from our benchmark set. Annotations with a depth of 3 make up almost half of all annotations in MFO before the removal (Figure S15b). After the removal, the most frequent annotations became of depth 5 (Figure S15a). In BPO, the most frequent annotations are of depth 5 or more, indicating a healthy increase of specific GO terms being added to our annotation database. In CCO, however, most new annotations in our benchmark set are of depth 3, 4 and 5 (Figure S15). This difference could partially explain why the same computational methods perform very differently in different ontologies, and benchmark sets. We have also calculated total information content per protein for the benchmark sets shown in Figure S16. Taxonomic distributions of the proteins in our final benchmark set are shown in Figure 14.

**Figure 14:**
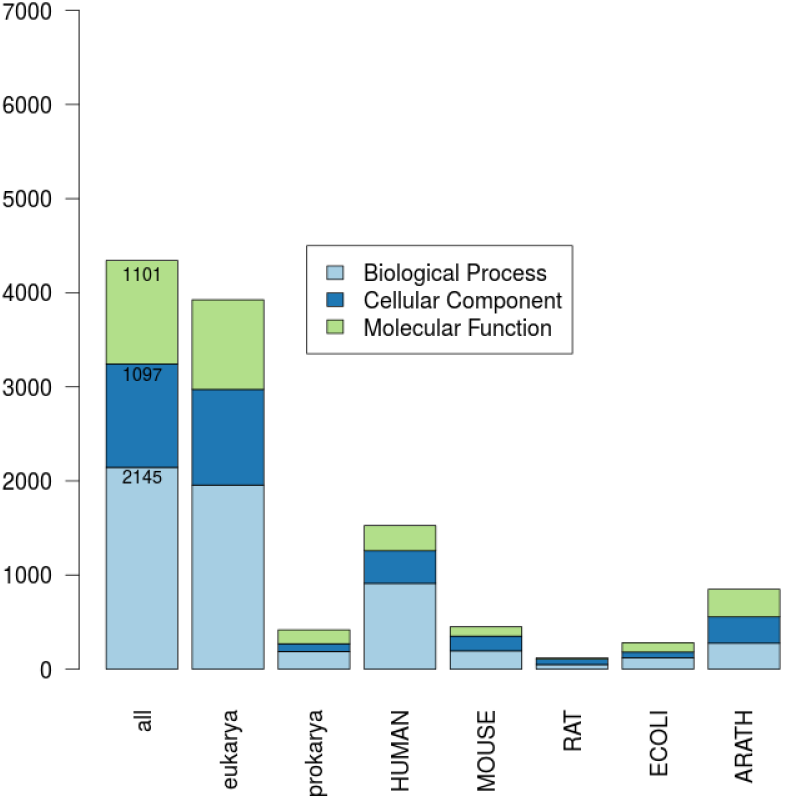
Number of proteins in each benchmark species and ontology.

Additional analyses were performed to assess the characteristics of the benchmark set, including the overall information content of the terms being annotated.

### 3.2 Protein-centric evaluation

Two main evaluation metrics were used in CAFA3, the *F*_max_ and the *S*_min_. The *F*_max_ based on the precision-recall curve, while the *S*_min_ is based the RU-MI curve. Mathematical definitions of these metrics are shown in pages 22 and 23 of Supplemental Materials. The RU-MI curve (51) takes into account the information content of each GO term in addition to counting the number of true positives, false positives, etc. See Supplemental Materials for their mathematical definitions. The information theory based evaluation metrics counters the high-throughput low-information annotations such as protein binding, but down-weighing these terms according to their information content, as the ability to predict such non-specific functions are not as desirable and useful and the ability to predict more specific functions.

The two assessment modes from CAFA2 were also used in CAFA3. In the partial mode, predictions were evaluated only on those benchmarks for which a model made at least one prediction. The full evaluation mode evaluates all benchmark proteins and methods were penalized for not making predictions. Evaluation results in Figures 3, 4, and 5 are made using the full evaluation mode. Evaluation results using the partial mode are shown in Figure S2 in the Supplemental Materials.

Two baseline models were also computed for these evaluations. The Naïve method assigns the term frequency as the prediction score for any protein, regardless of any protein-specific properties. BLAST was based on results using the Basic Local Alignment Search Tool (BLAST) software against the training database (52). A term will be predicted as the highest local alignment sequence identity among all BLAST hits annotated from the training database. Both of these methods were trained on the experimentally annotated proteins and their sequences in Swiss-Prot (53) at time *t*_0_.

### 3.3 Microbe screens

To assess matrix production, we used mutants from the PA14 NR collection (54). Mutants were transferred from the −80°C freezer stock using a sterile 48-pin multiprong device into 200μl LB in a 96-well plate. The cultures were incubated overnight at 37°C, and their OD600 was measured to assess growth. Mutants were then transferred to tryptone agar with 15g of tryptone and 15g of agar in 1L amended with Congo red (Aldrich, 860956) and Coomassie brilliant blue (J.T. Baker Chemical Co., F789-3). Plates were incubated at 37°C overnight followed by four day incubation at room temperature on allow the wrinkly phenotype to develop. Colonies were imaged and scored on Day 5. To assess motility, mutants were revived from freezer stocks as described above. After overnight growth, a sterile 48-pin multiprong transfer device with a pin diameter of 1.58 mm was used to stamp the mutants from the overnight plates into the center of swim agar made with M63 medium with 0.2% glucose and casamino acids and 0.3% agar). Care was taken to avoid touching the bottom of the plate. Swim plates were incubated at room temperature (19-22°C) for approximately 17 hours before imaging and scoring. Experimental procedures in *P. aeruginosa* to determine proteins that are associated with the two functions in CAFA-*π* are shown in Figure 15.

**Figure 15:**
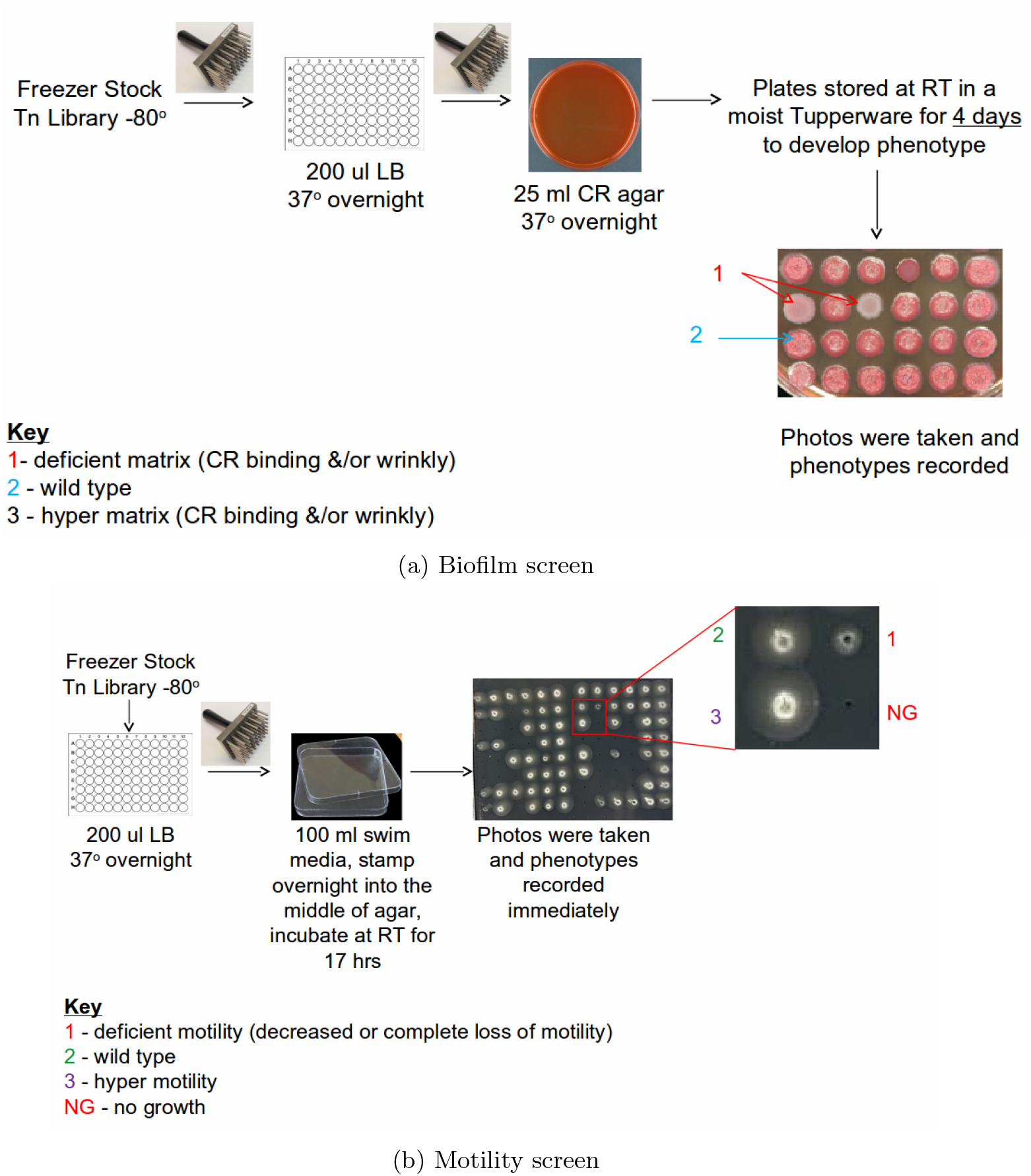
Experimental procedure of determining genes associated with the functions biofilm formation and motility in *P. aeruginosa*

Biofilm formation in *Candida albicans* was assessed in single gene mutants from the Noble (55) and GRACE (56) collections. In the Noble Collection, mutants of *C. albicans* have had both copies of the candidate gene deleted. Most of the mutants were created in biological duplicate. From this collection, 1274 strains corresponding to 653 unique genes were screened. The GRACE collection provided mutants with one copy of each gene deleted and the other copy placed under the control of a doxycycline-repressible promoter. To assay these strains, we used medium supplemented with 100μg/ml doxycycline strains, when rendered them functional null mutants. We screened 2348 mutants from the GRACE collection, 255 of which overlapped with mutants in the Noble collection, for 2746 total unique mutants screened in total. To assess defects in biofilm formation or biofilm-related traits, we performed two assays: (1) colony morphology on agar medium and (2) biofilm formation on a plastic surface (Figure 16). For both of these assays we used Spider medium, which was designed to induce hyphal growth in *C. albicans* (57), and which promotes biofilm formation (39). Strains were first replicated from frozen 96 well plates to YPD agar plates. Strains were then replicated from YPD agar to YPD broth, and grown overnight at 30°C. From YPD broth, strains were introduced onto Spider agar plates and into 96 well plates of Spider broth. When strains from the GRACE collection were assayed, 100μg/ml doxycycline was included in the agar and broth, and aluminium foil was used to protect the media from light. Spider agar plates inoculated with *C. albicans* mutants were incubated at 37°C for two days before colony morphologies were scored. Strains in Spider Broth were shaken at 225 rpm at 37°C for three days, and then assayed for biofilm formation at the air-liquid interface as follows. First, broth was removed by slowly tilting plates and pulling liquid away by running a gloved hand over the surface. Biofilms were stained by adding 100μl of 0.1 percent crystal violet dye in water to each well of the plate. After 15 minutes, plates were gently washed in three baths of water to remove dye without disturbing biofilms. To score biofilm formation for agar plates, colonies were scored by eye as either smooth, intermediate, or wrinkled. A wild-type colony would score wrinkled, and mutants with intermediate or smooth appearance were considered defective in colony biofilm formation. For biofilm formation on a plastic surface, the presence of a ring of cell material in the well indicated normal biofilm formation, while low or no ring formation mutants were considered defective. Genes whose mutations resulted defects in both or either assay were considered True for biofilm function. A complete list of the mutants identified in the screens is available in Table S1.

**Figure 16:**
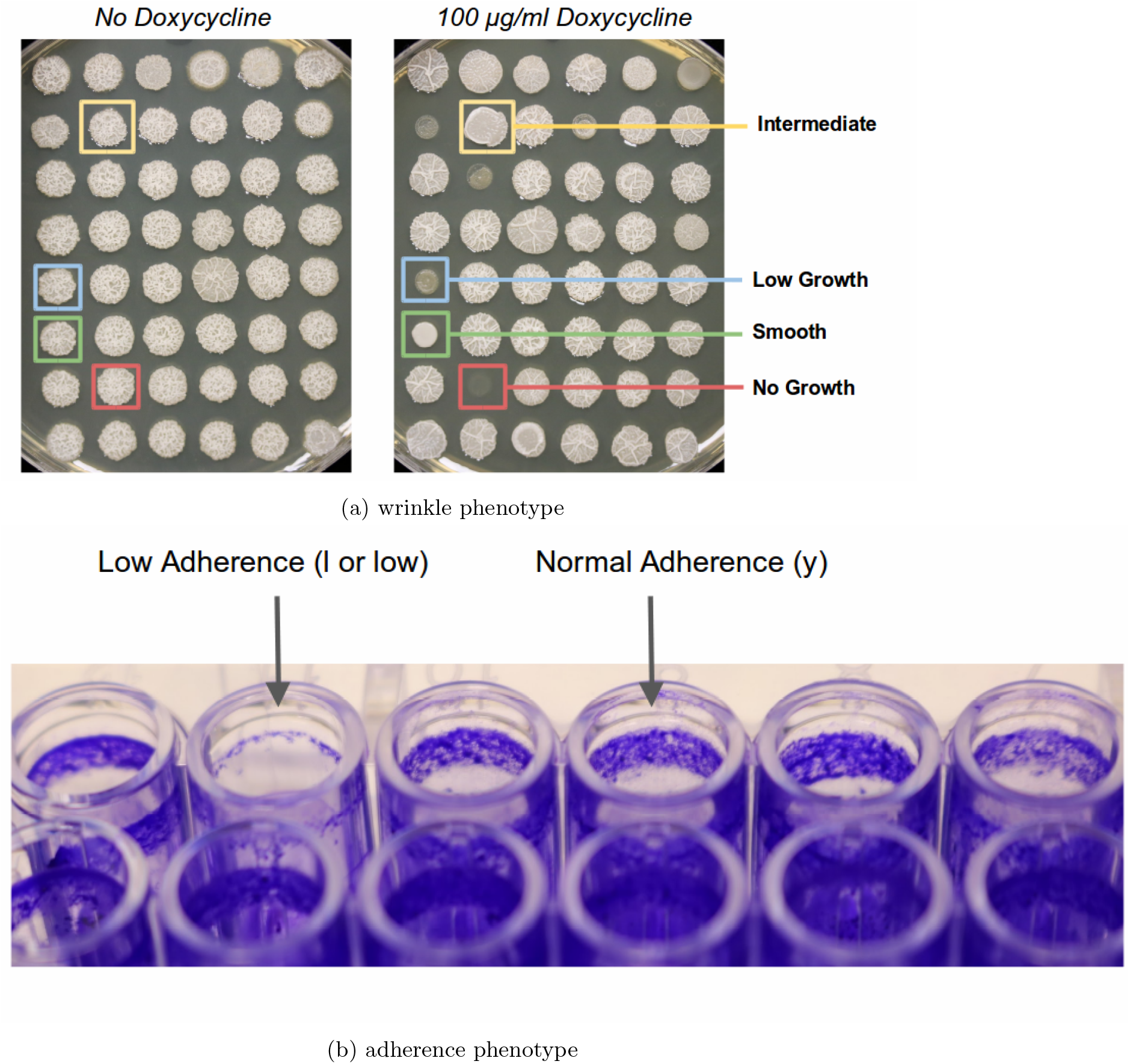
Experimental procedure of determining genes associated with the functions biofilm formation in *C. albicans*

A protein is considered True in the biofilm function, if its mutant phenotype is smooth or intermediate under Doxycycline.

### 3.4 Term-centric evaluation

The evaluations of the CAFA-*π* methods were based on the experimental results in Section 3.3. We adopted both *F*_max_ based on precision-recall curves and area under ROC curves. There are a total of six baseline methods, as described in Table 3.

**Table 3:** Baseline methods in term-centric evaluation of protein function prediction.

|  | Model Number | Training data | Score assignment |
| --- | --- | --- | --- |
| expression | 1 | Gene expression compendium for *P. aeruginosa* PAO1 | Highest correlation score out of all pairwise correlations |
|  | 2 |  | Top 10 average correlation score |
| blast | 1 | All experimental annotation in UniProt-GOA. Sequences from Swiss-Prot | Highest sequence identity out of all pairwise BLASTp hits |
|  | 2 | All experimental annotation in UniProt-GOA. Sequences from Swiss-Prot and TrEMBL |  |
| blastcomp | 1 | All experimental **and** computational annotations in UniProt-GOA. Sequences from Swiss-Prot |  |
|  | 2 | All experimental **and** computational annotations in UniProt-GOA. Sequences from Swiss-Prot and TrEMBL |  |

## 4 Discussion

Since 2010, the CAFA community has been a home to a growing group of scientists across the globe sharing the goal of improving computational function prediction. CAFA has been advancing this goal in three ways. First, through independent evaluation of computational methods against the set of benchmark proteins, thus providing a direct comparison of the methods’ reliability and performance at a given time point. Second, the challenge assesses the quality of the current state of the annotations, whether they are made computationally or not, and is set up to reliably track it over time. Finally, as described in this work, CAFA has started to drive the creation of new experimental annotations by facilitating synergies between different groups of researchers interested in function of biological macromolecules. These annotations not only represent new biological discoveries, but simultaneously serve to provide benchmark data for rigorous method evaluation.

CAFA3 and CAFA-*π* feature the latest advances in the CAFA series to create advanced and accurate methods for protein function prediction. We use the repeated nature of the CAFA project to identify certain trends via historical assessments. The analysis revealed that the performance of CAFA methods improved dramatically between CAFA1 and CAFA2. However, the protein-centric results for CAFA3 are mixed when compared to historical methods. Though the best performing CAFA3 method outperformed the top CAFA2 methods (Figure 1), this was not consistently true for other rankings. Among all three CAFA challenges, CAFA2 and CAFA3 methods inhabit the top 12 places in MFO and BPO. Between CAFA2 and CAFA3 the performance increase is more subtle. Based on the annotations of methods (Supplementary Materials), many of the top-ranking methods are improved versions of methods that have been evaluated in CAFA2. Interestingly, the top performing CAFA3 method, which consistently outperformed methods from all past CAFAs in the major categories, was a novel contribution (Zhu lab).

For this iteration of CAFA we performed genome-wide screens of phenotypes in *P. aeruginosa* and *C. albicans* as well as a targeted screen in *D. melanogaster*. This not only allowed us to assess the accuracy with which methods predict genes associated with select biological processes, but also to use CAFA as an additional driver for new biological discovery. In short, our experimental work identified more than a thousand of new functional annotations in three highly divergent species. Though all screens have certain limitations, the genome-wide screens also bypass questions of biases in curation. This evaluation provides key insights: CAFA3 methods did not generalize well to selected terms. Because of that, we ran a second effort, CAFA-*π*, in which participants focused solely on predicting the results of these targeted assays. This targeted effort led to improved performance, suggesting that when the goal is to identify genes associated with a specific phenotype, tuning methods may be required.

For CAFA evaluations, we have included both Naïve and sequence-based (BLAST) baseline methods. For the evaluation of *P. aeruginosa* screen results, we were also able to include a gene expression baseline from a previously published compendium (33). Intriguingly, the expression-based predictions outperformed existing methods for this task. In future CAFA efforts, we will include this type of baseline expression-based method across evaluations to continue to assess the extent to which this data modality informs gene function prediction. The results from the CAFA3 effort suggest that gene expression may be particularly important for successfully predicting term-centric biological process annotations.

The primary takeaways from CAFA3 are: (1) Genome-wide screens complement annotation-based efforts to provide a richer picture of protein function prediction; (2) The best performing method was a new method, instead of a light retooling of an existing approach; (3) Gene expression, and more broadly, systems data may provide key information to unlocking biological process predictions, and (4) Performance of the best methods has continued to improve. The results of the screens released as part of CAFA3 can lead to a re-examination of approaches which we hope will lead to improved performance in CAFA4.

## 5 Acknowledgements

Will be provided with the final manuscript

## 6 Data and Software

Data are available on figshare: https://figshare.com/articles/Supplementary_data/8135393 The assessment software used in this paper is available under GNU-GPLv3 license at: https://github.com/ashleyzhou972/CAFA_assessment_tool

## 7 Funding

The work of IF was funded, in part, by National Science Foundation award DBI-1458359. The work of CSG and AJL was funded, in part, by National Science Foundation award DBI-1458390 and GBMF 4552 from the Gordon and Betty Moore Foundation. The work of DAH and KAL was funded, in part, by National Science Foundation award DBI-1458390, National Institutes of Health NIGMS P20 GM113132, and the Cystic Fibrosis Foundation CFRDP STANTO19R0. The work of AP, HY, AR and MT was funded by BBSRC grants BB/K004131/1, BB/F00964X/1 and BB/M025047/1, Consejo Nacional de Ciencia y Tecnología Paraguay (CONACyT) grants 14-INV-088 and PINV15-315, and NSF Advances in Bio Informatics grant 1660648. DK acknowledges supports from the National Institutes of Health (R01GM123055) and the National Science Foundation (DMS1614777, CMMI1825941). PB acknowledges support from National Institutes of Health (R01GM60595). GB and BZK acknowledge support from the National Science Foundation (NSF 1458390) and NIH DP1MH110234. FS was funded by the ERC StG 757700 “HYPER-INSIGHT” and by the Spanish Ministry of Science, Innovation and Universities grant BFU2017-89833-P. FS further acknowledges funding from the Severo Ochoa award to the IRB Barcelona. The work of SK was funded by ATT Tieto käyttöön grant and Academy of Finland. TB and SM were funded by NIH awards UL1 TR002319 and U24 TR002306. The work of CZ and ZW was funded by National Institutes of Health R15GM120650 to ZW and start-up funding from the University of Miami to ZW. PR acknowledges NSF grant DBI-1458477. PT acknowledges support from Helsinki Institute for Life Sciences. The work of FZ and WT was funded by the National Natural Science Foundation of China (31671367, 31471245, 91631301) and the National Key Research and Development Program of China (2016YFC1000505, 2017YFC0908402]. CS acknowledges support by the Italian Ministry of Education, University and Research (MIUR) PRIN 2017 project 2017483NH8. SZ is supported by National Natural Science Foundation of China (No. 61872094 and No. 61572139) and Shanghai Municipal Science and Technology Major Project (No. 2017SHZDZX01). PLF and RLH were supported by the National Institutes of Health NIH R35-GM128637 and R00-GM097033. DTJ, CW, DC and RF were supported by the UK Biotechnology and Biological Sciences Research Council (BB/L020505/1 and BB/L002817/1) and Elsevier. The work of YZ and CZ was funded in part by the National Institutes of Health award GM083107, GM116960, AI134678, the National Science Foundation award DBI1564756, and the Extreme Science and Engineering Discovery Environment (XSEDE) award MCB160101 and MCB160124. The work of BG, VP, RD, NS and NV was funded by the Ministry of Education, Science and Technological Development of the Republic of Serbia, Project No. 173001. The work of YWL, WHL, JMC was funded by the Taiwan Ministry of Science and Technology (106-2221-E-004-011-MY2). YWL, WHL, JMC further acknowledge support from “the Human Project from Mind, Brain and Learning” of the NCCU Higher Education Sprout Project by the Taiwan Ministry of Education and the National Center for High-performance Computing for computer time and facilities.The work of IK and AB was funded by Montana State University and NSF Advances in Biological Informatics program through grant number 0965768. BR, TG and JR are supported by the Bavarian Ministry for Education through funding to the TUM. The work of RB, VG, MB, and DCEK was supported by the Simons Foundation and NIH NINDS grant number 1R21NS103831-01.

## Notes

https://figshare.com/articles/Supplementary_data/8135393

https://github.405com/ashleyzhou972/CAFA_assessment_tool

