## Supplementary files for "The CAFA challenge reports improved protein function prediction and new functional annotations for hundreds of genes through experimental screens"

### List of Tables

S1 Number of experimental annotations in UniProt-GOA for biofilm formation (GO:0042710) . . 17
S2 Number of experimental annotations in UniProt-GOA for cilium or flagellum-dependent cell motility (GO:0001539) . . . . . . . . . . . . . . . . . . . . . . . . . . . . . . . . . . . . . 18
S3 Number of experimental annotations in UniProt-GOA for long-term memory (GO:0007616) . 19

### Contents

**Mathematical definitions of protein-centric metrics** **22**

**List of CAFA3 Keywords** **24**

Figure S1: Head-to-head comparison between top five CAFA3 versus CAFA1 methods

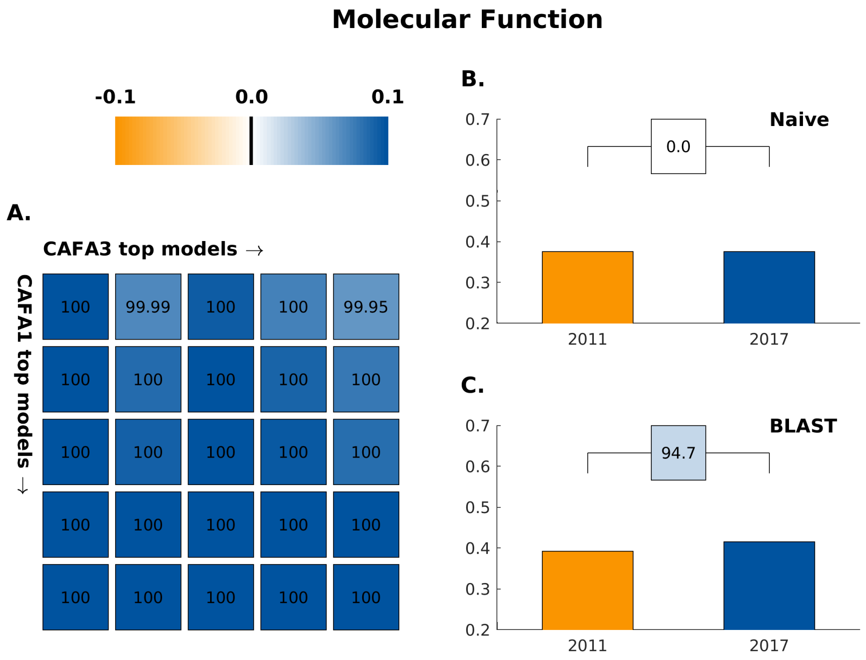

(a)

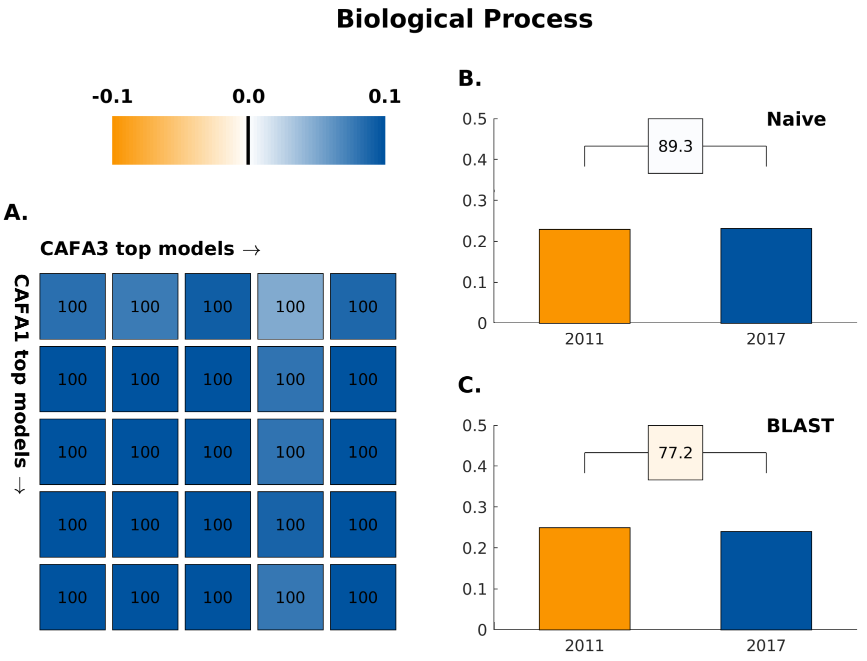

(b)

Figure S2: $F_{\max}$ curves for the top-performing methods in *partial* evaluation mode for (A) Molecular Function ontology, (B) Biological Process ontology, (C) Cellular Component ontology on *No Knowledge* benchmark.

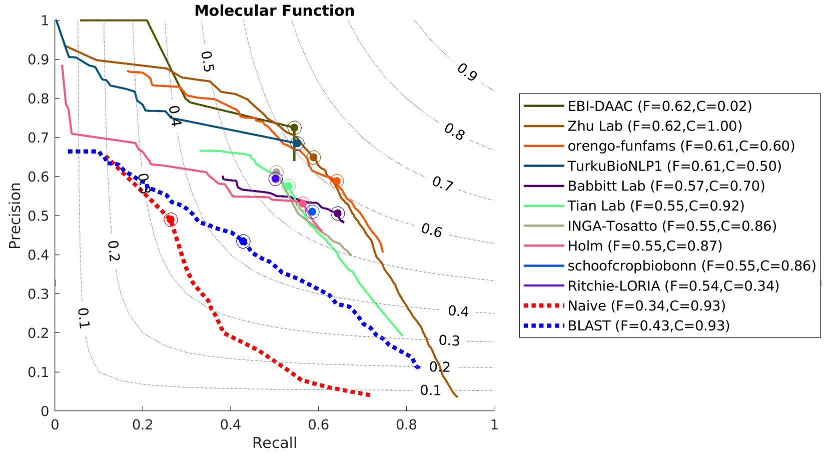

(a)

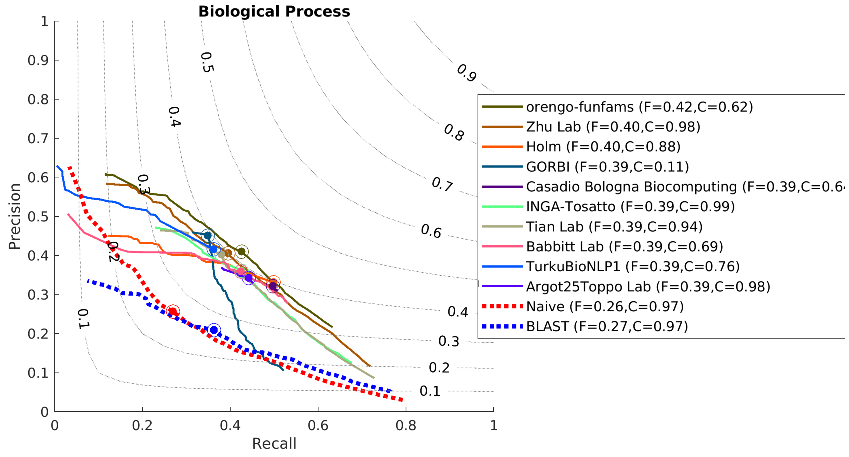

(b)

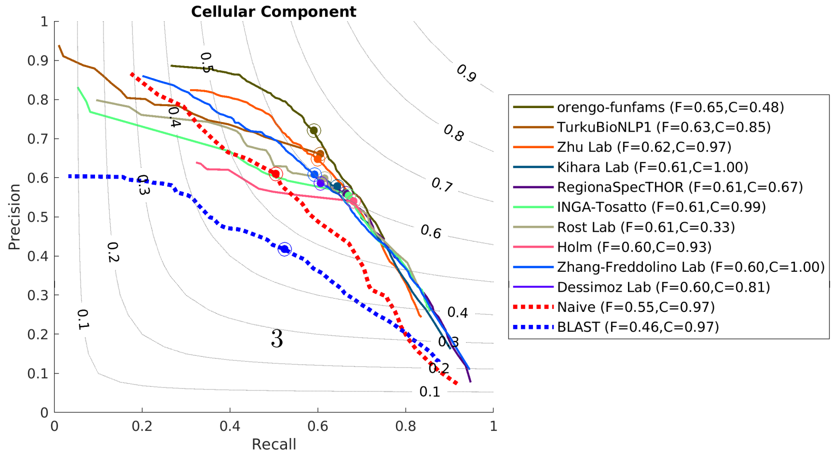

(c)

Figure S3: $F_{\max}$ curves for the top-performing methods on *limited knowledge* benchmarks for (A) Molecular Function ontology, (B) Biological Process ontology, (C) Cellular Component ontology in *full* evaluation mode

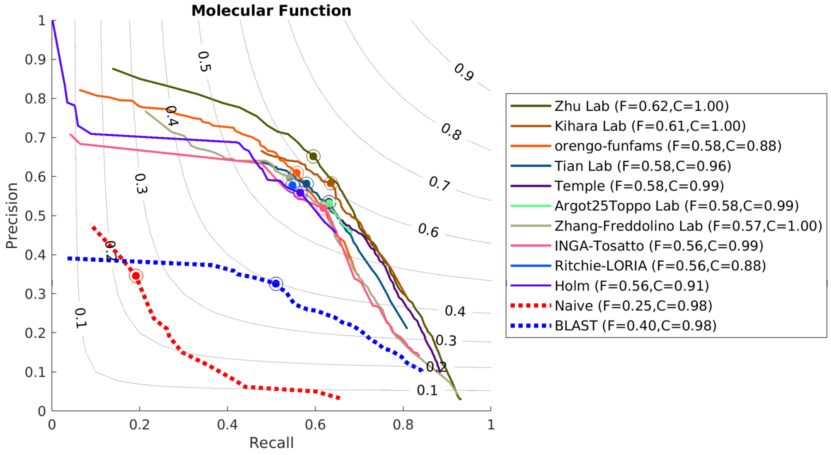

(a)

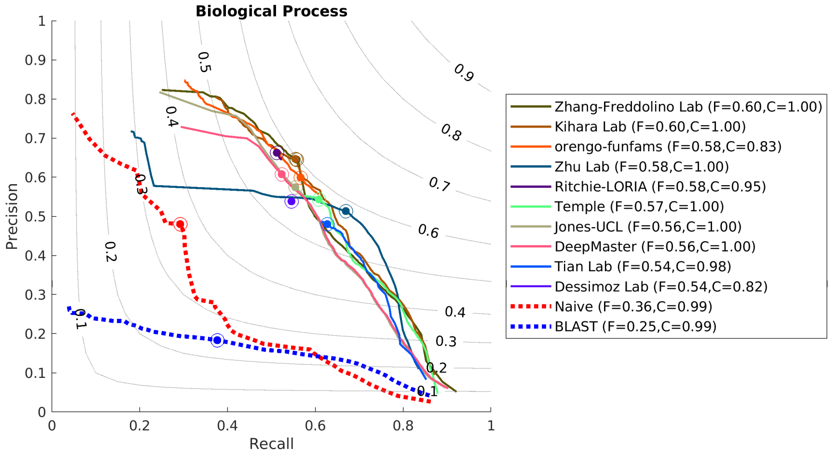

(b)

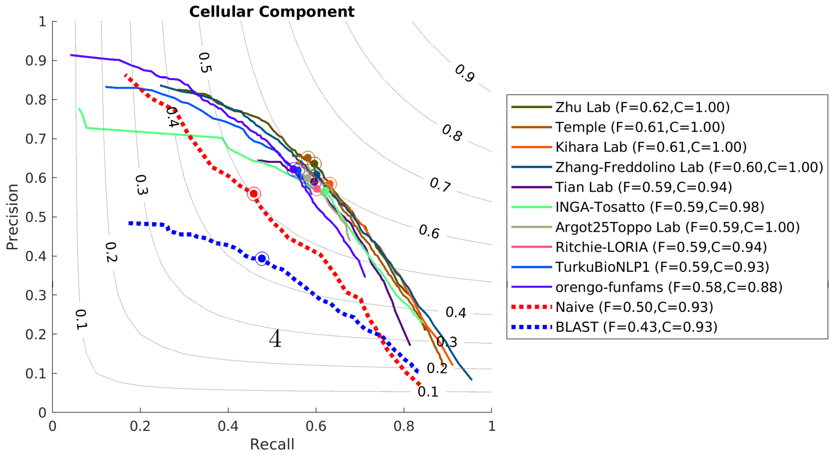

(c)

Figure S4: Weighted precision-recall curves for the top-performing methods for (A) Molecular Function ontology, (B) Biological Process ontology, (C) Cellular Component ontology on *No Knowledge* benchmark and *full* evaluation mode

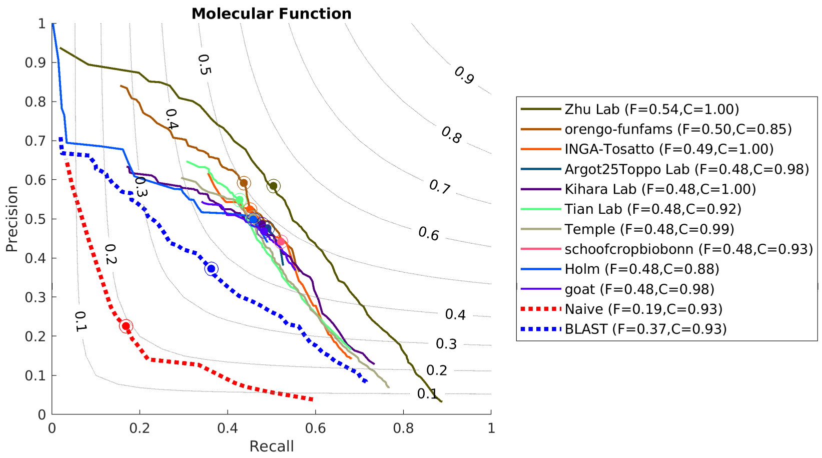

(a)

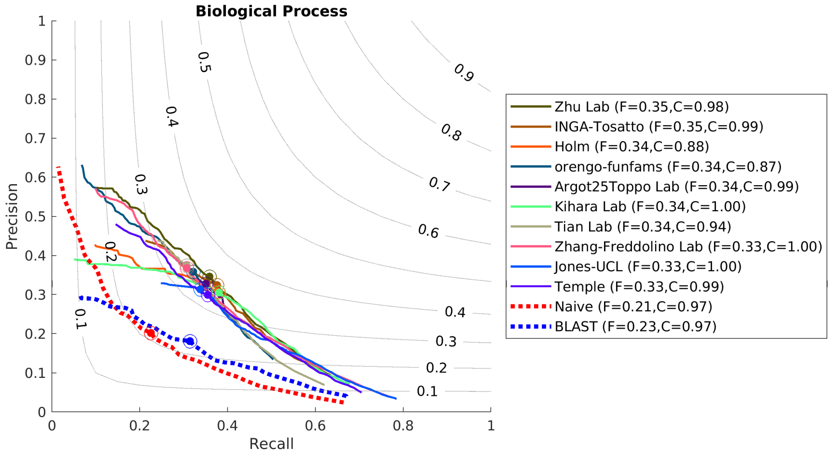

(b)

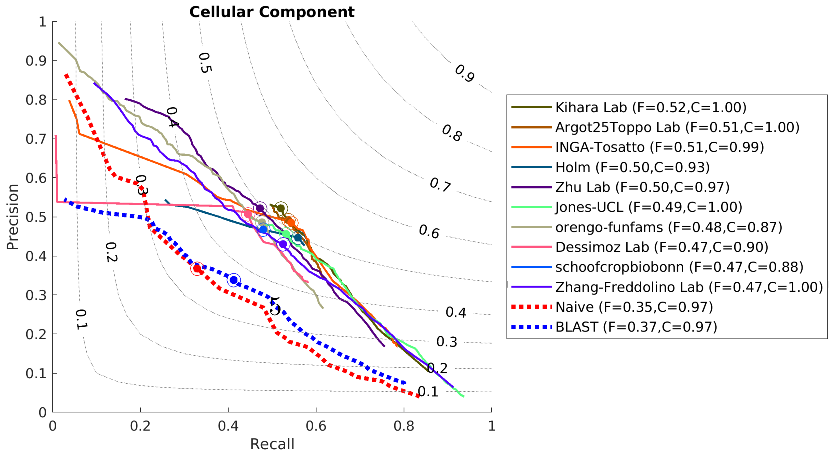

(c)

Figure S5: Normalized RU-MI curves for the top-performing methods for (A) Molecular Function ontology, (B) Biological Process ontology, (C) Cellular Component ontology on *No Knowledge* benchmark and *full* evaluation mode

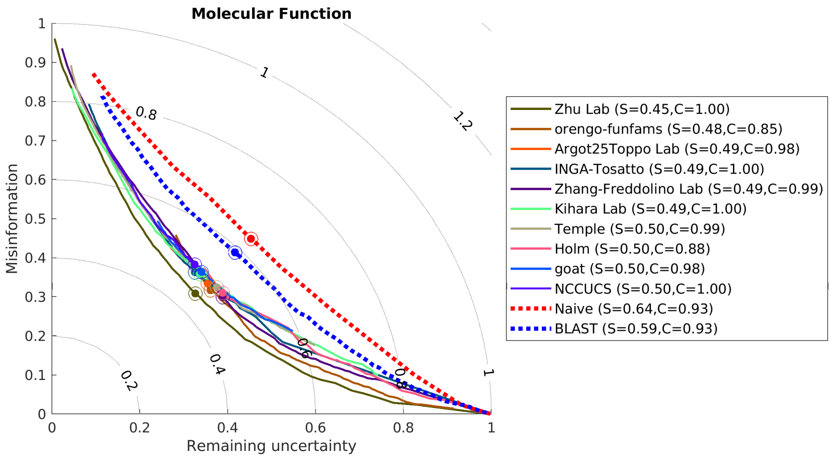

(a)

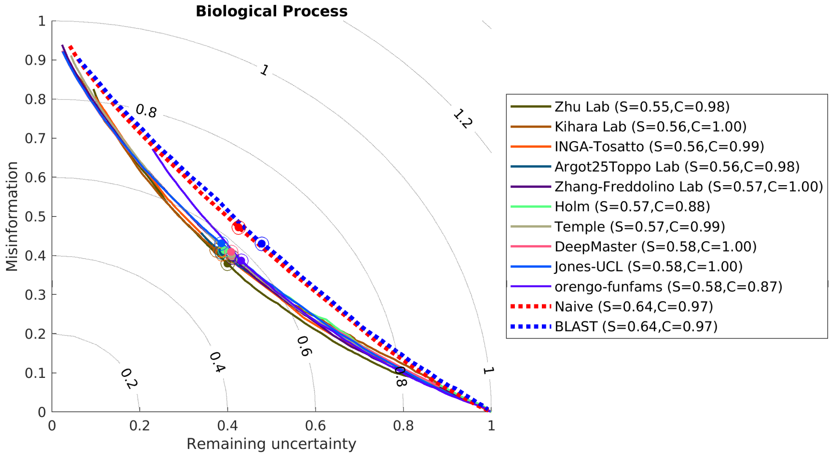

(b)

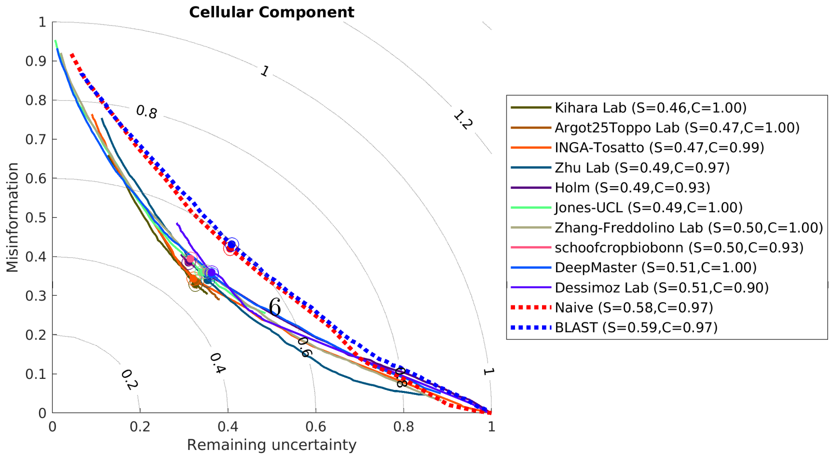

(c)

The followings figures S6, S7, S8, S9, S10, S11, S12, S13, S14 shows evaluations based on the *No Knowledge* benchmarks in the *full* evaluation mode.

Figure S6: Top 10 $F_{\max}$ in *Homo Sapiens*

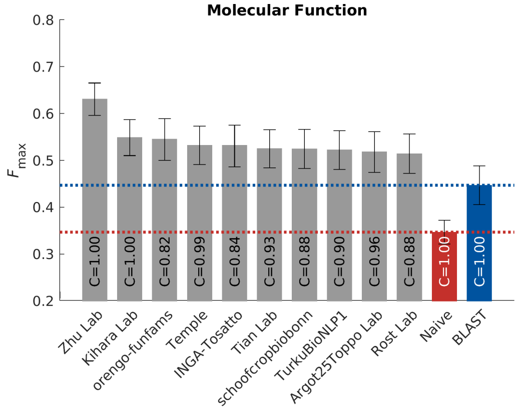

(a)

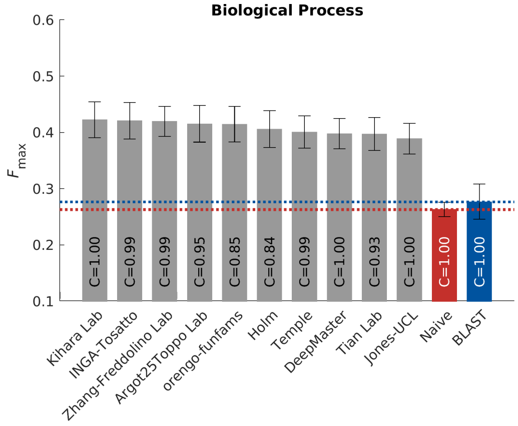

(b)

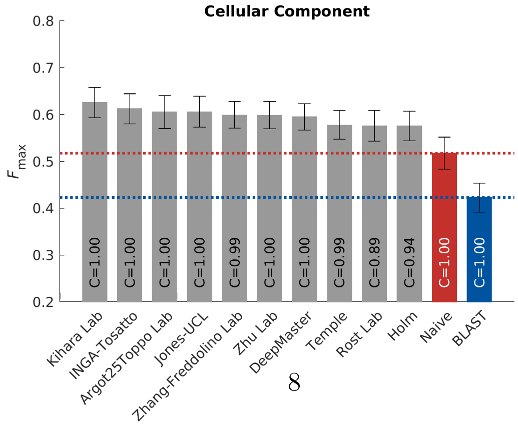

(c)

#### Figure S7: Top 10 $F_{\max}$ in *Arabidopsis thaliana*

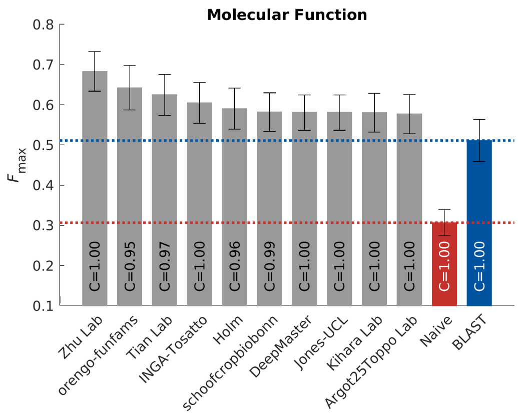

(a)

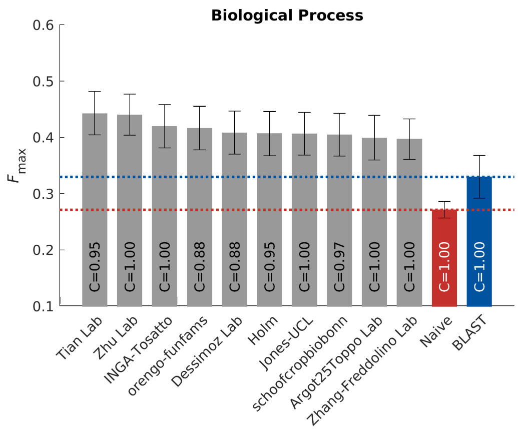

(b)

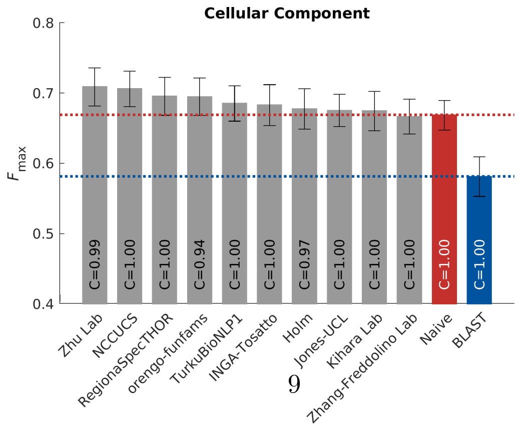

(c)

Figure S8: Top 10 $F_{\max}$ in *Mus musculus*

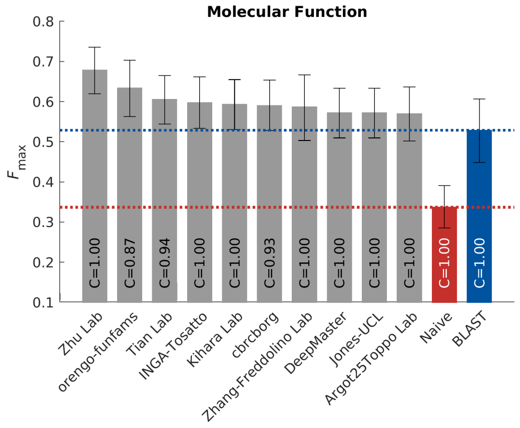

(a)

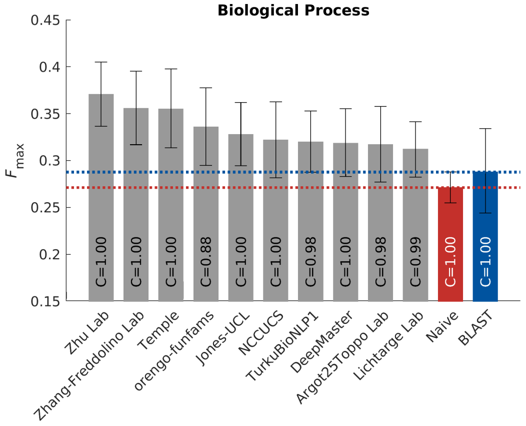

(b)

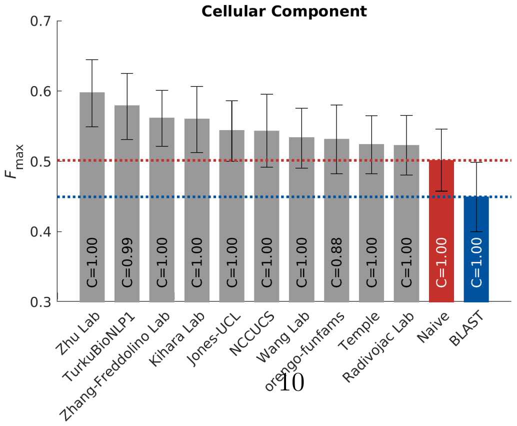

(c)

Figure S9: Top 10 $F_{\max}$ in *Rattus norvegicus*

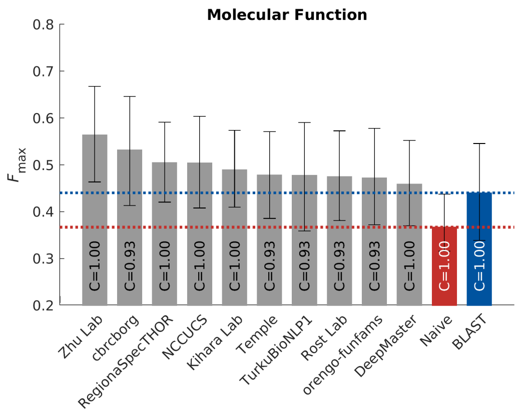

(a)

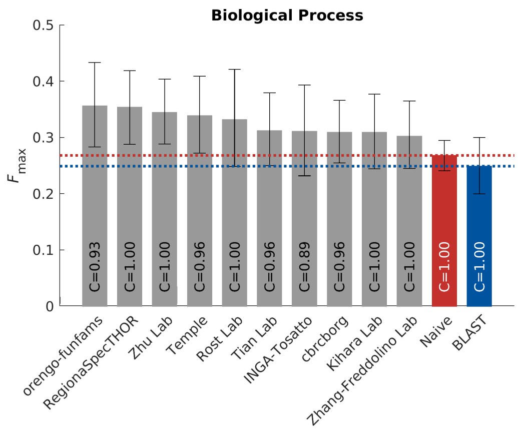

(b)

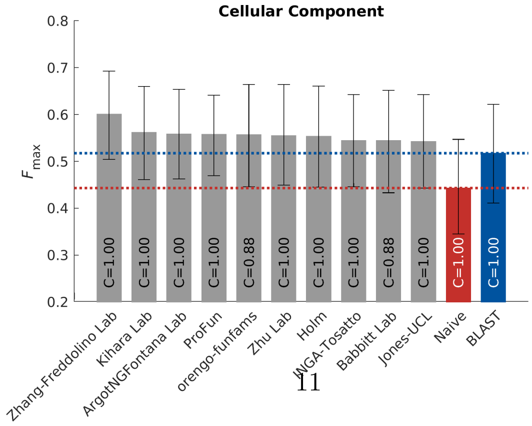

(c)

Figure S10: Top 10 $F_{\max}$ in *Escherichia coli K12*

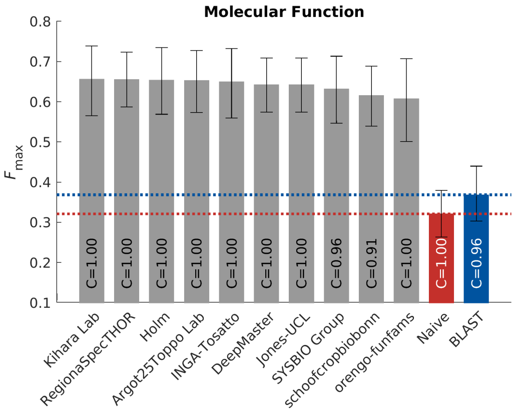

(a)

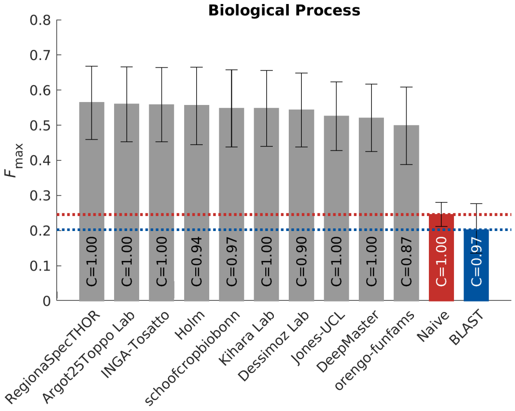

(b)

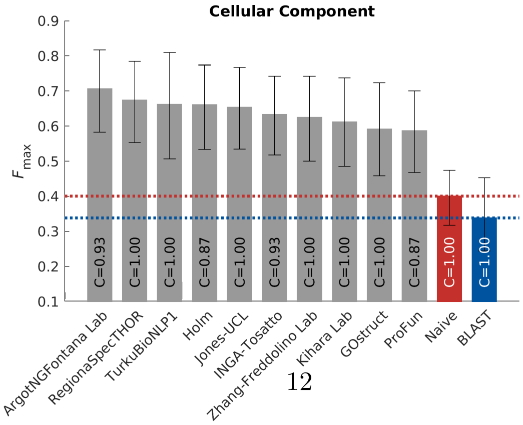

(c)

Figure S11: Top 10 $F_{\max}$ in *Drosophila melanogaster*

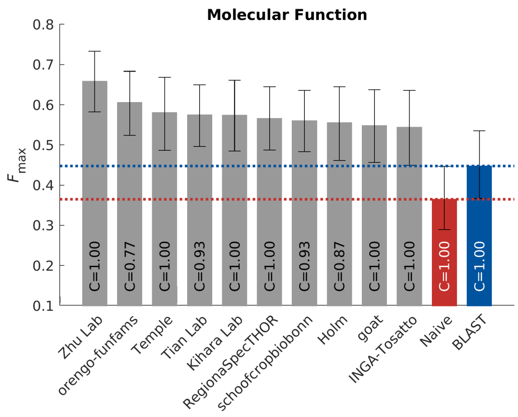

(a)

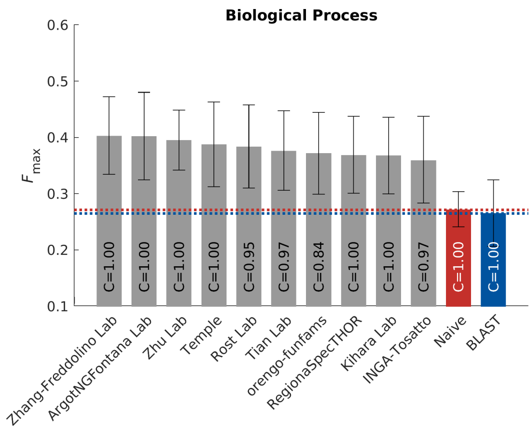

(b)

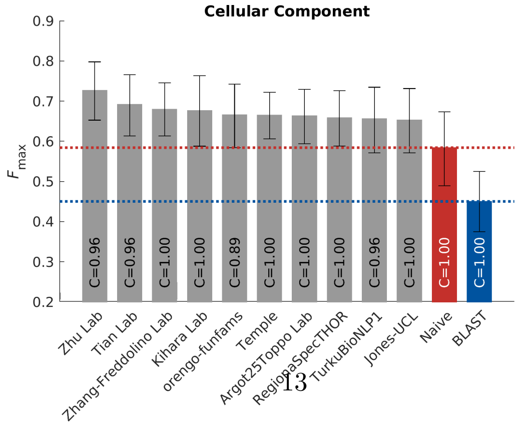

(c)

Figure S12: Top 10 $F_{\max}$ in *Dictyostelium discoideum*

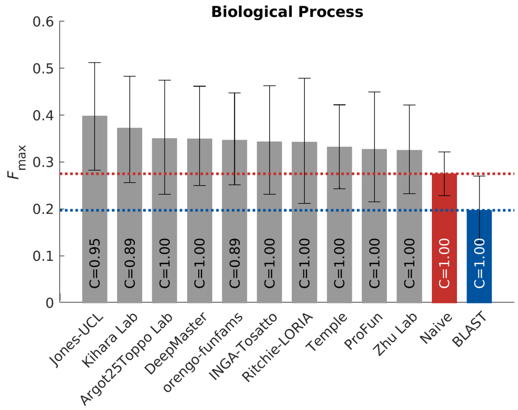

(a)

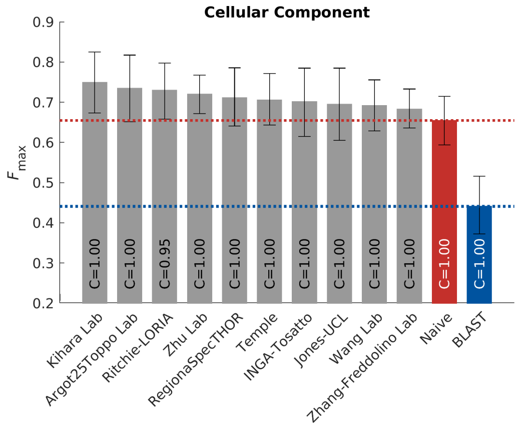

(b)

Figure S13: Top 10 $F_{\max}$ in *Danio rerio*

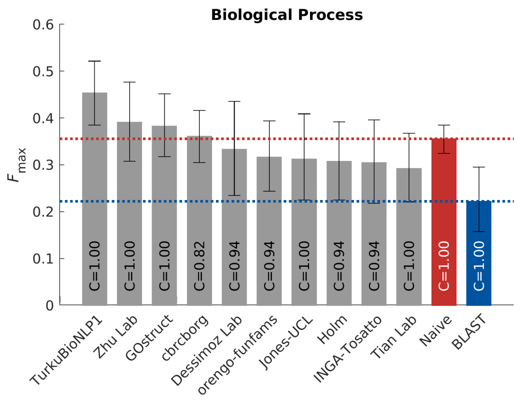

(a)

Figure S14: Top 10 $F_{\max}$ in *Candida albicans (strain SC5314 / ATCC MYA-2876)*

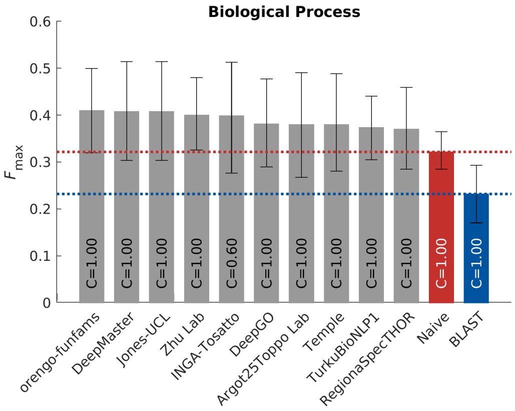

(a)

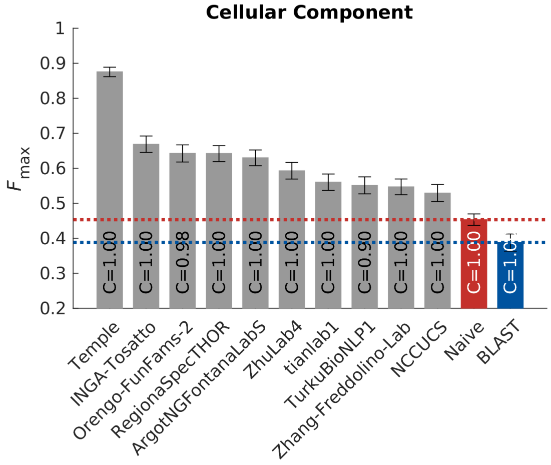

(b)

Table S1: Number of experimental annotations in UniProt-GOA for biofilm formation (GO:0042710)

| Taxonomy ID | Scientific Name | Number of experimental annotations |
|---|---|---|
| 237561 | Candida albicans SC5314 | 188 |
| 208964 | Pseudomonas aeruginosa PAO1 | 54 |
| 83333 | Escherichia coli K-12 | 46 |
| 559292 | Saccharomyces cerevisiae S288C | 10 |
| 284593 | [Candida] glabrata CBS 138 | 5 |
| 578454 | Candida parapsilosis CDC317 | 4 |
| 190486 | Xanthomonas citri pv. citri str. 306 | 2 |
| 290339 | Cronobacter sakazakii ATCC BAA-894 | 2 |
| 216592 | Escherichia coli 042 | 2 |
| 5476 | Candida albicans | 2 |
| 224308 | Bacillus subtilis subsp. subtilis str. 168 | 1 |
| 1314 | Streptococcus pyogenes | 1 |
| 330879 | Aspergillus fumigatus Af293 | 1 |
| 210007 | Streptococcus mutans UA159 | 1 |
| 243277 | Vibrio cholerae O1 biovar El Tor str. N16961 | 1 |
| 227321 | Aspergillus nidulans FGSC A4 | 1 |
| 93061 | Staphylococcus aureus subsp. aureus NCTC 8325 | 1 |
| 316273 | Xanthomonas campestris pv. vesicatoria str. 85-10 | 1 |
| 1280 | Staphylococcus aureus | 1 |
| 100226 | Streptomyces coelicolor A3(2) | 1 |

Table S2: Number of experimental annotations in UniProt-GOA for cilium or flagellum-dependent cell motility (GO:0001539)

| Taxonomy ID | Scientific Name | Number of experimental annotations |
|---|---|---|
| 208964 | Pseudomonas aeruginosa PAO1 | 41 |
| 3055 | Chlamydomonas reinhardtii | 40 |
| 83333 | Escherichia coli K-12 | 30 |
| 185431 | Trypanosoma brucei brucei TREU927 | 26 |
| 7227 | Drosophila melanogaster | 16 |
| 10090 | Mus musculus | 7 |
| 7955 | Danio rerio | 5 |
| 9606 | Homo sapiens | 4 |
| 287 | Pseudomonas aeruginosa | 4 |
| 85962 | Helicobacter pylori 26695 | 3 |
| 224308 | Bacillus subtilis subsp. subtilis str. 168 | 2 |
| 189518 | Leptospira interrogans serovar Lai str. 56601 | 1 |
| 5664 | Leishmania major | 1 |
| 246197 | Myxococcus xanthus DK 1622 | 1 |
| 9913 | Bos taurus | 1 |
| 529507 | Proteus mirabilis HI4320 | 1 |
| 9615 | Canis lupus familiaris | 1 |
| 99287 | Salmonella enterica subsp. enterica serovar Typhimurium str. LT2 | 1 |
| 8090 | Oryzias latipes | 1 |
| 31286 | Trypanosoma brucei rhodesiense | 1 |

Table S3: Number of experimental annotations in UniProt-GOA for long-term memory (GO:0007616)

| Taxonomy ID | Scientific Name | Number of experimental annotations |
|---|---|---|
| 7227 | Drosophila melanogaster | 217 |
| 10090 | Mus musculus | 23 |
| 10116 | Rattus norvegicus | 22 |
| 6239 | Caenorhabditis elegans | 13 |
| 9606 | Homo sapiens | 7 |
| 381128 | Lehmannia marginata | 1 |

Figure S15: Distribution of benchmark depth in leaf nodes. A leaf node is defined if any descendent nodes are not included as benchmark.

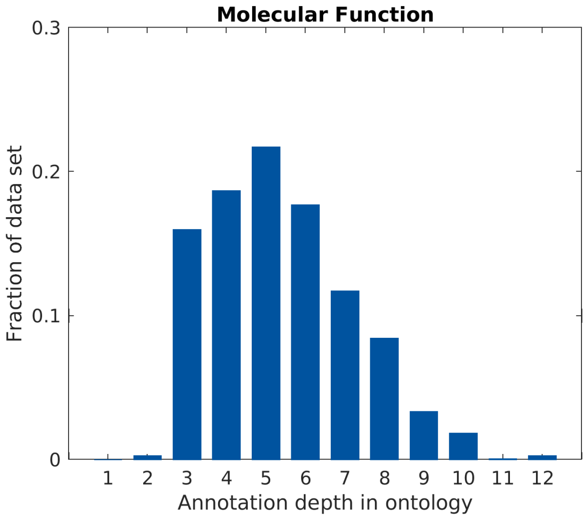

(a) After removing single protein-binding annotations

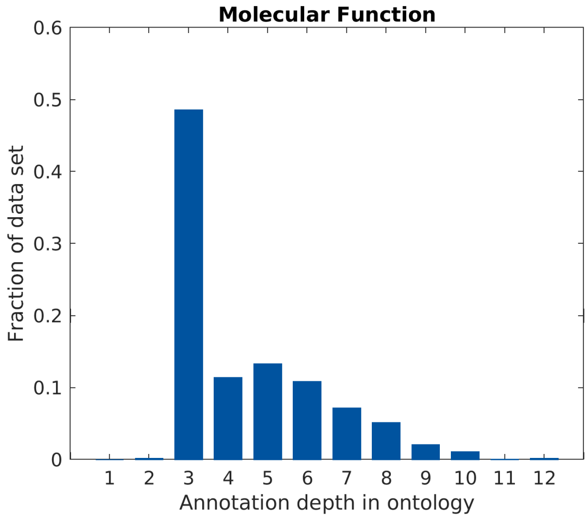

(b) Before removing single protein-binding annotations

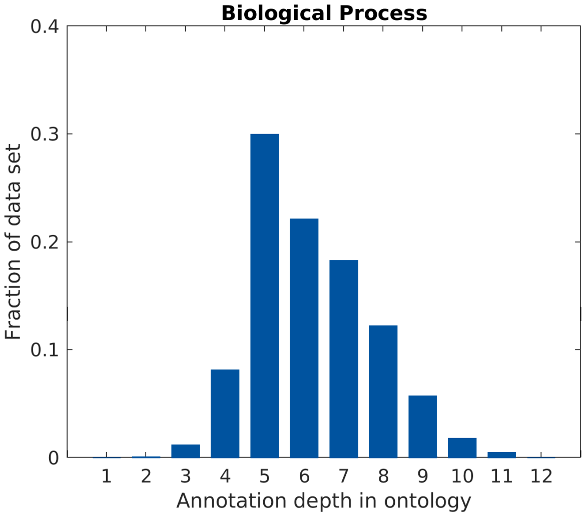

(c)

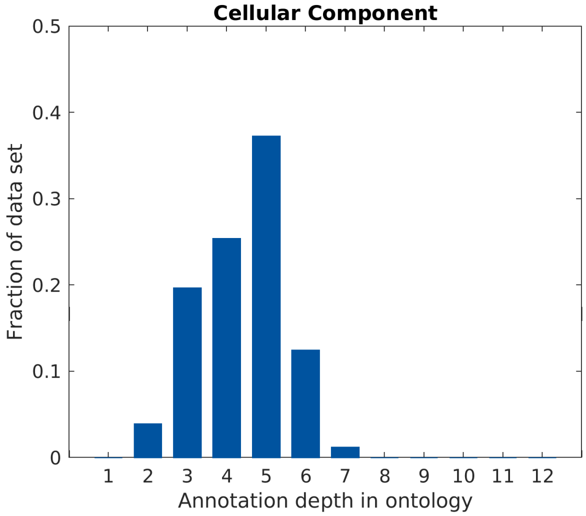

(d)

Figure S16: Frequency of total information content of benchmark proteins for (a) Molecular Function ontology, (b) Biological Process ontology, and (c) Cellular Component ontology. Data include all benchmark proteins and all experimentally annotated proteins at the point of benchmark collection $t_1$. The red point indicates the value of information content for the predicted annotation using to the Naïve model.

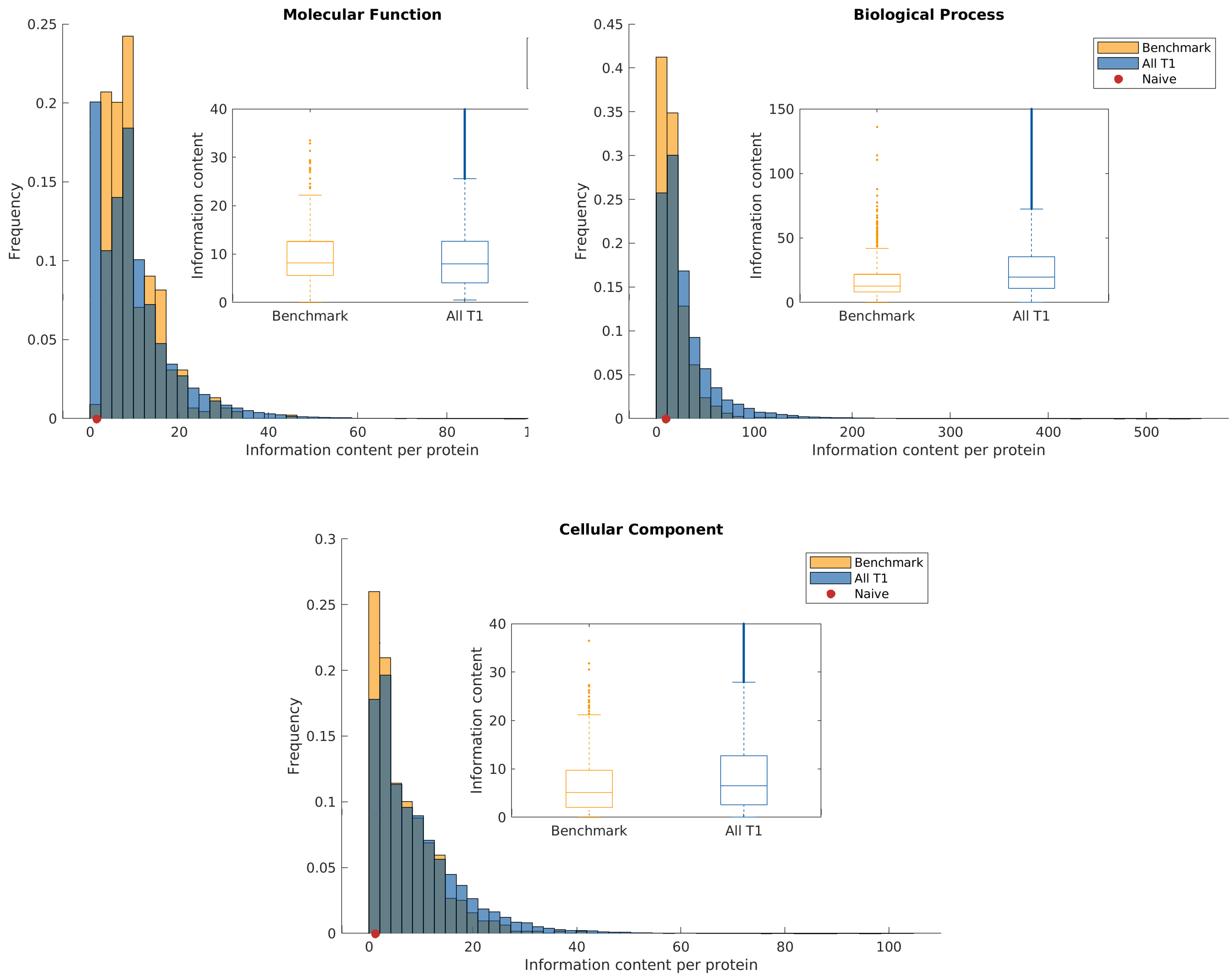

#### Mathematical definitions of protein-centric metrics

- Precision Recall

$$
\begin{aligned}
pr(\tau) &= \frac{1}{m(\tau)} \sum_{i=1}^{m(\tau)} \frac{\sum_f \mathbb{1}\left(f \in P_i(\tau) \wedge f \in T_i\right)}{\sum_f \mathbb{1}\left(f \in P_i(\tau)\right)}, \\
rc(\tau) &= \frac{1}{n_e} \sum_{i=1}^{n_e} \frac{\sum_f \mathbb{1}\left(f \in P_i(\tau) \wedge f \in T_i\right)}{\sum_f \mathbb{1}\left(f \in T_i\right)}, \\
F_{\max} &= \max_{\tau} \left\{ \frac{2 \cdot pr(\tau) \cdot rc(\tau)}{pr(\tau) + rc(\tau)} \right\},
\end{aligned}
$$

where $P_i(\tau)$ denotes the set of terms that have predicted scores greater than or equal to $\tau$ for a protein sequence $i$, $T_i$ denotes the corresponding ground-truth set of terms for that sequence, $m(\tau)$ is the number of sequences with at least one predicted score greater than or equal to $\tau$, $\mathbb{1}(\cdot)$ is an indicator function and $n_e$ is the number of targets used in a particular mode of evaluation. In the full evaluation mode $n_e = n$, the number of benchmark proteins, whereas in the partial evaluation mode $n_e = m(0)$; i.e., the number of proteins which were chosen to be predicted using the particular method. For each method, we refer to $\frac{m(0)}{n}$ as the *coverage* because it provides the fraction of benchmark proteins on which the method made any predictions.

- Remaining Uncertainty and Missing Information

$$
\begin{aligned}
ru(\tau) &= \frac{1}{n_e} \sum_{i=1}^{n_e} \sum_f ic(f) \cdot \mathbb{1}\left(f \notin P_i(\tau) \wedge f \in T_i\right), \\
mi(\tau) &= \frac{1}{n_e} \sum_{i=1}^{n_e} \sum_f ic(f) \cdot \mathbb{1}\left(f \in P_i(\tau) \wedge f \notin T_i\right), \\
S_{\min} &= \min_{\tau} \left\{ \sqrt{ru(\tau)^2 + mi(\tau)^2} \right\},
\end{aligned}
$$

where $ic(f)$ is the information content of the ontology term $f$ [**?**]. It is estimated in a maximum likelihood manner as the negative binary logarithm of the conditional probability that the term $f$ is present in a protein's annotation given that all its parent terms are also present. Note that here, $n_e = n$ in the full evaluation mode and $n_e = m(0)$ in the partial evaluation mode applies to both $ru$ and $mi$.

- Weighted Precision Recall

$$\begin{aligned} wpr(\tau) &= \frac{1}{m(\tau)} \sum_{i=1}^{m(\tau)} \frac{\sum_f ic(f) \cdot \mathbb{1}\left(f \in P_i(\tau) \wedge T_i(\tau)\right)}{\sum_f ic(f) \cdot \mathbb{1}\left(f \in P_i(\tau)\right)}, \quad \text{and} \\ wrc(\tau) &= \frac{1}{n_e} \sum_{i=1}^{n_e} \frac{\sum_f ic(f) \cdot \mathbb{1}\left(f \in P_i(\tau) \wedge T_i(\tau)\right)}{\sum_f ic(f) \cdot \mathbb{1}\left(f \in T_i(\tau)\right)}, \end{aligned}$$

where $P_i(\tau)$ is the set of predicted terms for protein $i$ with score no less than threshold $\tau$ and $T_i$ is the set of true terms for protein $i$, $m(\tau)$ is the number of sequences with at least one predicted score greater than or equal to $\tau$, and $n_e$ is the number of proteins used in a particular mode of evaluation. In the full evaluation mode $n_e = n$, the number of benchmark proteins, whereas in the partial evaluation mode $n_e = m(0)$.

- Normalized Remaining Uncertainty and Missing Information

$$\begin{aligned} nru(\tau) &= \frac{1}{n_e} \sum_{i=1}^{n_e} \frac{\sum_f ic(f) \cdot \mathbb{1}\left(f \notin P_i(\tau) \wedge f \in T_i\right)}{\sum_f ic(f) \cdot \mathbb{1}\left(f \in P_i(\tau) \vee f \in T_i\right)}, \quad \text{and} \\ nmi(\tau) &= \frac{1}{n_e} \sum_{i=1}^{n_e} \frac{\sum_f ic(f) \cdot \mathbb{1}\left(f \in P_i(\tau) \wedge f \notin T_i\right)}{\sum_f ic(f) \cdot \mathbb{1}\left(f \in P_i(\tau) \vee f \in T_i\right)}, \end{aligned}$$

where $P_i(\tau)$ is the set of predicted terms for protein $i$ with score no less than threshold $\tau$ and $T_i$ is the set of true terms for protein $i$, and $n_e$ is the number of proteins used in a particular mode of evaluation. In the full evaluation mode $n_e = n$, the number of benchmark proteins, whereas in the partial evaluation mode $n_e$ is the number of proteins that have at least one positive predicted score.

#### List of CAFA3 Keywords

sequence alignment, sequence-profile alignment, profile-profile alignment, phylogeny, sequence properties, physicochemical properties, predicted properties, protein interactions, gene expression, mass spectrometry, genetic interactions, protein structure, literature, genomic context, synteny, structure alignment, comparative model, predicted protein structure, de novo prediction, machine learning, genome environment, operon, ortholog, paralog, homolog, hidden Markov model, clinical data, genetic data, natural language processing, other functional information
